# Reconstitution of multistep recruitment of ULK1 to membranes in autophagy

**DOI:** 10.1101/2025.11.07.687251

**Authors:** Yongjia Duan, Ye Lu, Sanjoy Paul, Johannes Betz, Léa P Wilhelm, Annan S. I. Cook, Xuefeng Ren, Elias Adriaenssens, Sascha Martens, Ian G. Ganley, Gerhard Hummer, James H. Hurley

**Author notes:** These authors contributed equally (Yongjia Duan, Ye Lu, Sanjoy Paul, Johannes Betz, Léa P Wilhelm, Annan S. I. Cook). Corresponding authors: IGG,; GHH,; JHH. **Teaser:** The ULK1 complex is targeted to membranes by a WF motif of ATG101, ATG13-WIPI binding, and an ULK1 PVP motif that docks on ATG13.

## Abstract

The Human ULK1 autophagy-initiating complex consists of ULK1, FIP200, and the HORMA domain heterodimer ATG13:ATG101. PI3P is essential to recruit ULK1C to membranes for ULK1, but ULK1C subunits do not contain PI3P-binding domains. Here we show that the ATG13:ATG101 dimer forms a complex with the PI3P-binding protein WIPI3, as well as WIPI2. Bound to WIPI2-3, ATG13:ATG101 inserts its Trp-Phe (WF) finger into the membrane. MD simulations show that WIPIs and the WF finger cooperatively stabilizes the complex on membranes. Biochemical reconstitution and cell-based assays show that WIPI3:ATG13 engagement promotes ATG16L1 phosphorylation, autophagy and mitophagy. A kinase domain (KD)-proximal PVP motif in the ULK1 IDR docks onto the ATG13:ATG101 HORMA dimer brings the ULK1 KD close to the membrane. The PVP motif is essential for *in vitro* ULK1 phosphorylation of ATG16L1 and important for autophagy and mitophagy. These data establish a stepwise pathway for recruitment of the ULK1 KD to the vicinity of the membrane surface.

## Introduction

Macroautophagy (henceforth autophagy) is a conserved bulk degradative process that maintains cellular homeostasis under stresses such as starvation, mitochondrial damage, abnormal protein aggregation, and invasion by intracellular pathogens(*1–3*). The dysfunction of autophagy is implicated in neurodegenerative disease, with particularly strong genetic evidence linking defective autophagy to Parkinson’s disease. Autophagy is initiated by the co-recruitment and activation of the Unc51-like kinase 1 complex (ULK1C) and the class III phosphatidylinositol-3 kinase complex I (PI3KC3-C1)(*4, 5*). The co-recruitment and mutual stabilization of these two complexes, together with small vesicles containing the lipid scramblase ATG9, is decisive in determining whether or not autophagy progresses.

The human ULK1 complex consists of the ULK1 kinase itself, the large scaffold protein FIP200, and the two HORMA (Hop/Rev7/Mad2) domain regulatory subunits ATG13 and ATG101(*4, 6–9*). ATG13 is conserved and essential for autophagy throughout all eukaryotes. In fission yeast, metazoa, and much of biology, ATG13 forms a 1:1 heterodimer with another essential autophagy factor, ATG101. The heterodimer is mediated by the HORMA domains of these proteins, which forms a stable and extensive structural interface. ATG101 contains a protruding Trp-Phe (WF) finger motif, which is important for autophagy initiation(*10–12*), but whose precise function is unknown. ATG101 consists of barely more than a single HORMA domain, with the sole addition of an amphipathic C-terminal helix (CTH) immediately following the HORMA domain. As for ATG13, after the N-terminal HORMA domain, the rest of ATG13 consists of a ∼250 residues intrinsically disordered region (IDR). The most C-terminal part of the ATG13 IDR is responsible for binding to FIP200 and ULK1(*13, 14*). The ATG13:ATG101 HORMA dimer contains a binding site for ATG9, which presumably serves to promote the interaction between ULK1C and ATG9 vesicles. Yet mutational disruption of ATG13:ATG9 binding only partially blocks autophagy(*15*), therefore, it seems that ATG13 and ATG101 must have functions beyond ATG9 binding.

The class III phosphatidylinositol (PI) 3-kinase complex I (PI3KC3-C1) is responsible for the conversion of PI to PI 3-phosphate (PI3P), a critical early step in the progression of autophagy. Mammalian PI3KC3-C1 consists of the lipid kinase VPS34, the pseudokinase VPS15, and the regulatory subunits BECN1 and ATG14(*4, 16*). The production of PI3P in autophagic membranes recruits the WD40 interacting with phosphoinositide (WIPI) proteins 1-4, which in turn recruit the bridge-like lipid transport (BLTP) ATG2 and the ATG12-ATG5-ATG16L1 complex. ATG2 drives membrane expansion by transferring phospholipids from the endoplasmic reticulum (ER) to the growing phagophore(*17*). The ATG12-ATG5-ATG16L1 complex serves as a ubiquitin E3-like factor to promote the covalent conjugation of ATG8 proteins, which in humans consist of LC3A-C, GABARAP, and GABARAPL1-2, to the membrane lipid phosphatidylethanolamine (PE). PI3P-bound WIPI2 is critical for the recruitment of ATG12-ATG5-ATG16L1 downstream of PI3KC3-C1 activation(*18*). Many of the proteins in these pathways, including all the subunits of PI3KC3-C1, WIPI2, and ATG16L1, are substrates of ULK1 phosphorylation(*19–23*).

ULK1C and PI3KC3-C1 function is central to autophagy initiation and therefore closely coordinated at multiple levels. ULK1C and PI3KC3-C1 can form a physical supercomplex(*13*). ULK1 directly phosphorylates all four of the subunits of PI3KC3-C1. Moreover, ULK1C recruitment as judged by formation of ULK1 or ATG13 puncta is dependent upon the activity of PI3KC3(*18*). The size of ULK1 nanoclusters, and the average number of ULK1 molecules at the autophagosome is sensitive to PI3P synthesis. Inhibition of PI3KC3 activity reduces recruitment of ULK1 and ATG13 to the phagophore(*24, 25*), thus reducing autophagic flux. PI3P sensing was initially proposed to be mediated by a cluster of basic residues in the N-terminus of ATG13, on the basis of dot-blotting with immobilized membrane-free PI3P monomers(*26*). However, the crystal structure of the human ATG13:ATG101 HORMA dimer did not show evidence of a PI3P-binding pocket(*11*), nor, as detailed below, has it been possible to replicate the finding of PI3P binding using more stringent and biologically realistic giant unilamellar vesicle (GUV) binding assays. Thus, it is not known how PI3P production stabilizes ULK1C recruitment. Indeed, the mechanism for regulated translocation of ULK1C to phospholipid membranes upon autophagy induction, one of the most central events of autophagy initiation, has been entirely unresolved.

Recently, it was demonstrated that in the context of BNIP3/NIX-dependent mitophagy, the recruitment of key early autophagy initiating proteins proceeds in a non-standard order. BNIP3/NIX recruits WIPI2, which in turn recruits ULK1C via an interaction with a short linear sequence motif of ATG13 located just C-terminal to the HORMA domain(*27, 28*). We reasoned that this or other ATG13 interactions with WIPIs might provide the missing link to explain PI3P-dependent recruitment and stabilization of ULK1C on membranes. Here, we show that engagement of the ATG13:ATG101 HORMA dimer with WIPIs is a critical driver of ULK1C recruitment downstream of PI3KC3-C1 activity.

A further conundrum in the recruitment and activation of ULK1 in autophagy stems from the domain structure of ULK1 and its mode of integration in the full ULK1C. The protein kinase domain (KD) of ULK1 resides in the N-terminal part of the protein. The KD is connected by a 500-amino acid IDR to a C-terminal tandem MIT domain, which is also known as the early autophagy targeting (EAT) domain(*19*). The EAT domain is responsible for directly engaging with the rest of the ordered core of the ULK1C, which consists of the N-terminal 640 residues of FIP200 and the C-terminal part of the ATG13 IDR. Thus, the catalytic “business end” of ULK1 is separated by the membrane recruitment unit by 500 residues of IDR within ULK1 itself and an additional ∼150 residues of IDR within ATG13. We reasoned there must be additional mechanisms to bring the ULK1 kinase closer to the membrane, where most of its autophagic substrates are localized. Here, we show that a region of the ULK1 IDR proximal to the KD contains a short linear interaction motif that binds to the ATG13:ATG101 HORMA, so delivering the kinase domain to its site of action at the membrane surface.

## Results

### ATG13 contains a DHF motif that binds to WIPI3 at a different site than WIPI2

We took inspiration from the recent discovery that the HORMA-proximal portion of the ATG13 IDR contains a WIPI2 interacting region (W2IR)(*27*) that functions in BNIP3/NIX mitophagy. We sought to determine if WIPI-ULK1C interactions could serve more generally in PI3P-dependent membrane targeting of ULK1C by screening for interactions with other members of the WIPI family. We generated high-quality models of ATG13:ATG101 and WIPI3 by combining DeepMSA2 and AlphaFold2. We first used the DeepMSA2 server to generate multiple sequence alignments for downstream structure predictions and evolutionary analyses(*29*). Inspection of the aligned sequences of ATG13 revealed a highly conserved motif in the ATG13 IDR including residues Asp 213, His 214 and Phe 215 (hereafter, the “DHF motif“) (Fig. 1A, 1B). We noticed the signature sequence of “β-strand-(X)n-Φ-X-X-Φ-X-X-X-Ψ-F” (Φ is a hydrophobic residue and Ψ is an aromatic residue) in this motif, which is similar to the WIPI binding region (WIR) of ATG2A(*30*) (Fig. S1A, S1B), but divergent from the ATG16 W2IR motif (Fig. S1C)(*30, 31*). Because of this sequence similarity, we hypothesized that the predicted WIR of ATG13 might be the critical region binding to WIPI3.

**Figure 1.**
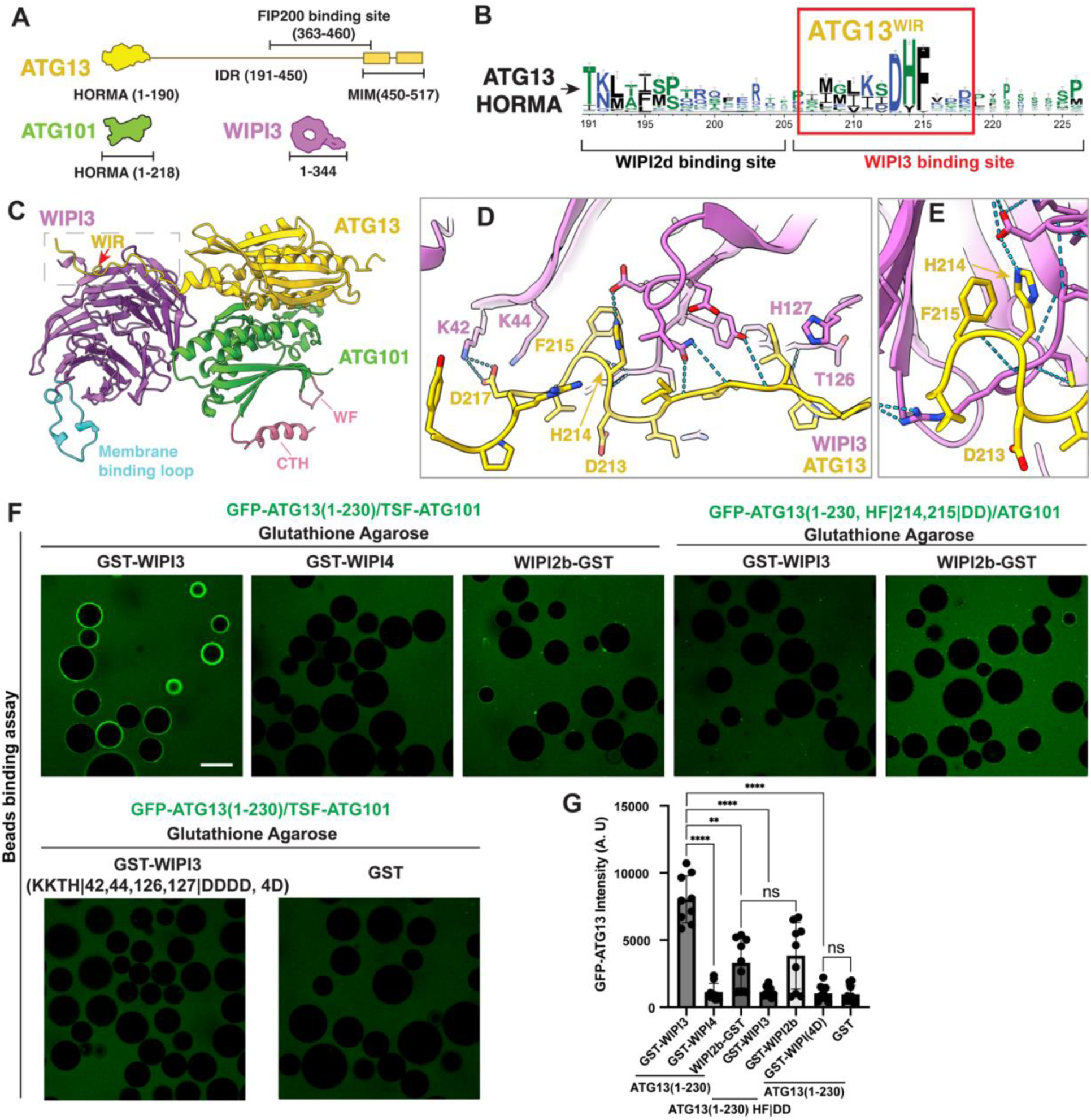
Identification of a WIPI3 interacting region in ATG13. **(A)** Schematic showing the HORMA, IDR, MIM domain and FIP200 binding site on ATG13, sequence length of ATG101 and WIPI3. **(B)** Bioinformatic analysis of the WIR motif of ATG13. Conservation in bits shown on the Y-axis with a red box showing the conserved fragment that mediates WIPI3 binding. **(C)** Structural model of the ATG13(1-220)-ATG101-WIPI3 tripartite complex predicted using AlphaFold2. **(D)** Close up view of the interface between the ATG13 W3IR and WIPI3 K42, K44, H127 and T126. **(E)** ATG13-D214 and H215 form hydrogen bonds to WIPI3 which are shown by the dashed line. **(F)** GST beads binding assay confirmed the strong and specific binding between ATG13 DHF motif with WIPI3-K42, K44, H127 and T126. All scale bars equal 100 µm. **(G)** Quantification analysis of GST beads binding assay, statistics: One-way ANOVA + Bonferroni post-hoc test. Three biological replicates were performed, with three images taken for each replicate. The average fluorescence intensity of all beads in each image was calculated. A.U represents arbitrary units. **P < 0.01, ****P < 0.0001 and ns: not significant. All data are mean ± s.d.

To address this possibility, we used the ColabFold implementation of AlphaFold2 to predict the structure of ATG13 1-220 in a complex with WIPI3 (Fig. 1C, 1D, S1D and S1E). We analyzed the predicted complex structure and indeed found that the His and Phe residues of the conserved DHF motif (residues 213-215) of ATG13 were positioned in the WIR-binding pocket of WIPI3 (Fig. 1C, 1D). Moreover, Lys42 and Lys44 of WIPI3 and Asp 217 (downstream of the DHF motif) of ATG13 are close enough to form salt bridges (Fig. 1D, 1E). To validate the AlphaFold predicted model, we performed a microscopy-based protein–protein interaction assay between ATG13:ATG101 and WIPI protein immobilized on GST-agarose beads. The results shows that there was a clear preference for WIPI3 over WIPI4 binding (Fig.1F, 1G). The double mutation of the ATG13 DHF motif (H214D and F215D, hereafter ATG13(HF|DD)) nearly abrogated the interaction between ATG13:ATG101 and WIPI3 (Fig. 1F, 1G), consistent with the structural hypothesis that the His-Phe dyad drives the interaction between ATG13 and WIPI3.

To test the contribution of these residues in WIPI3 to the interactions with the ATG13:ATG101 heterodimer, we produced the mutants of GST-WIPI3(K42D, K44D, T126D and H127D, abbreviated as WIPI3(4D)) and measured the binding with GFP-ATG13(1-230):ATG101. The 4D mutant showed a profound reduction in binding affinity (Fig. 1F, 1G). We sought to compare this interaction to the recently described between ATG13 and WIPI2(*27*). As a control, we also performed a GST beads binding assay comparing the binding of ATG13(1-230)/ATG101 to WIPI2 and WIPI3. The results confirmed the interaction between ATG13:ATG101 and WIPI2 as in a previous study(*27*) in the case where the HORMA is included (Fig 1F, 1G). Furthermore, mutating the ATG13 WIR motif at residues H214 and F215 did not substantially affect the WIPI2:ATG13:ATG101 interaction (Fig. 1F, 1G). This is consistent with the structural prediction and indicates that the interaction between WIPI2 and ATG13:ATG101 does not involve the ATG13 WIR motif. Taken together, these observations support a model for ATG13:ATG101 binding to WIPI3 that is driven by the WIR motif in the IDR of ATG13, which is C-terminal to the previously characterized WIPI2d binding site.

### ATG101 binds directly to phospholipid membrane through its WF-finger and C-terminal helix

We next considered how the assembly of the ATG13:ATG101 HORMA heterodimer with WIPI3 might dock onto membranes. WIPI3 is a member of the PROPPIN family of proteins, which dock onto membranes via a bispecific PI3P and PI3,5P_2_-binding (F/L)RRG motif and an adjacent hydrophobic membrane-binding loop(*32, 33*). We noted a coplanar orientation of the WF-finger and C-terminal helix (CTH) of ATG101 with the hydrophobic membrane binding loop of WIPI3 (Fig. 1C). Mutation of the ATG101 WF and CTH have demonstrated autophagy phenotypes(*10, 12*), yet their biochemical roles and interactors have been unknown. We wondered if these motifs might bind directly to the lipid bilayer and thereby recruit the ULK1 complex to PI3P-positive membranes. To test this hypothesis, we prepared GUVs containing 5% PI3P (Fig. 2A). Upon the addition of 200 nM WIPI3 to the reaction, we saw robust membrane recruitment of ATG13(1-230)-GFP:ATG101 to GUVs (Fig. 2B, 2C). We introduced mutations of ATG101 WF finger Trp110, Phe112 to Asp (ATG101(WF|DD)), and produced a CTH-truncated ATG101 (ATG101^ΔCTH^) and a construct combining the WF|DD and CTH truncation (ATG101^ΔCTH^ (WF|DD)). The loss of the CTH showed a quantifiable defect in membrane recruitment, while conversion of WF to DD more substantially diminished membrane binding (Fig. 2D, 2E). The double mutant construct showed almost the same effect with the WF to DD mutant, indicating that the WF finger makes a major contribution to membrane binding (Fig. 2D, 2E). In the presence of both WIPI3 and PI3P, the recruitment of the ATG13:101 WF mutant to GUV membranes showed a linear, concentration-dependent relationship (Fig. 2D, 2E). These data suggest when the WF finger is lost, residual membrane binding is primarily driven by the WIPI3-PI3P interaction. However, the ATG101 WF finger plays a critical role in facilitating the efficient recruitment of ATG13:ATG101 to the membrane at the physiologically relevant concentration of 40-50 nM(*34*).

**Figure 2.**
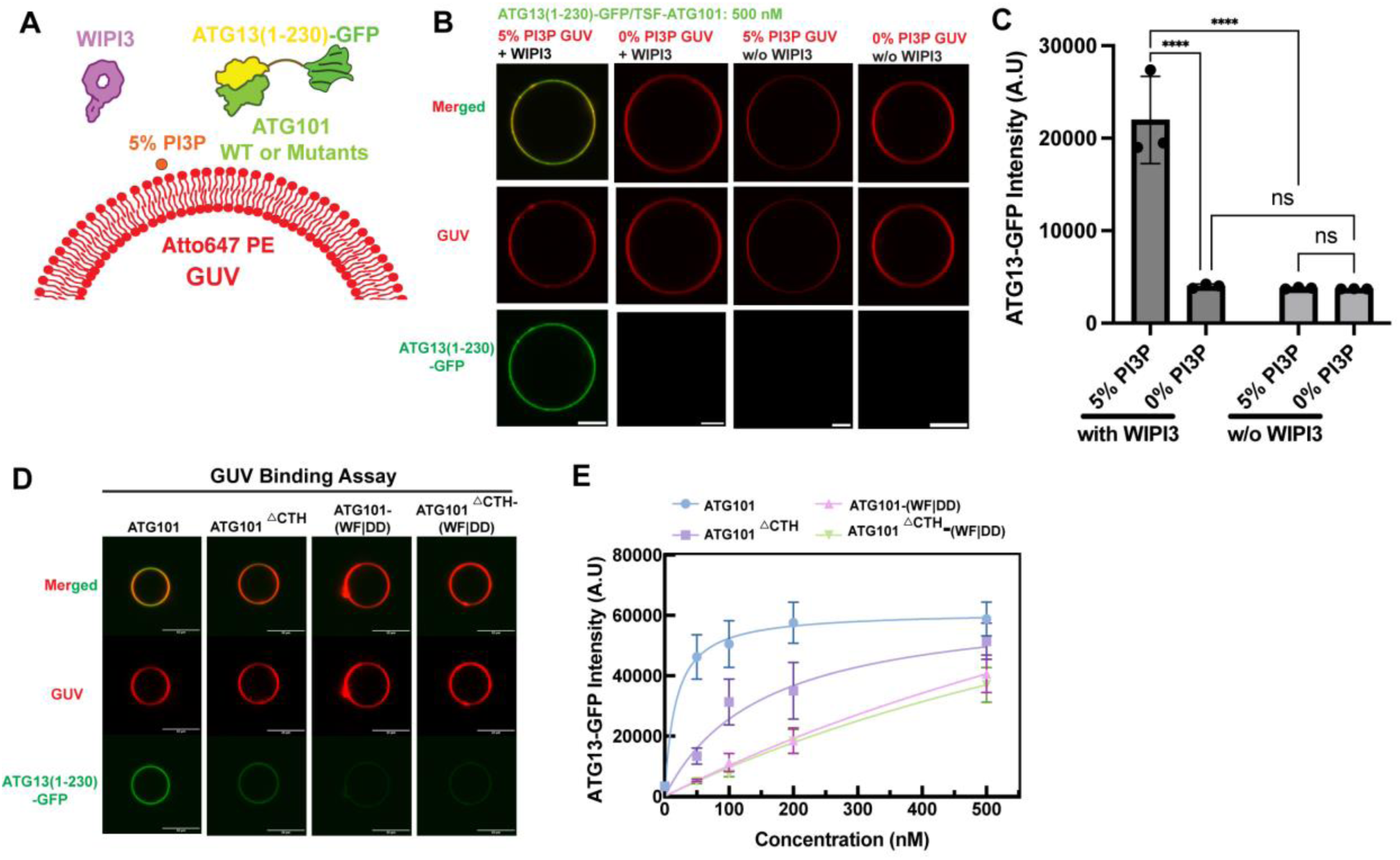
WF finger and the C-terminal helix of ATG101 are required for membrane binding. **(A)** Schematic of the GUV-based experiment. The GUVs which were rendered in red in the figure, were prepared with 5% PI3P and 0.1% Atto647 PE. GFP tagged ATG13(1-230):ATG101 was included at variable concentrations, while WIPI3 was included at 200 nM. **(B)** Confocal fluorescence images of GUV recruitment of ATG13(1-230)-GFP:ATG101 (500 nM) under the condition of with or without 5% PI3P and WIPI3 (200 nM). GUV recruitment of ATG13(1-230):ATG101 relies on WIPI3 and PI3P. The scale bar is 5 µm. **(C)** Quantitation of fluorescent intensity of ATG13(1-230):ATG101 binding on GUV at different condition as in (B). Three biological replicates were performed, at least three GUVs for each replicate were calculated. Statistics: Two-way ANOVA + Tukey’s multiple comparisons test. **(D)** Confocal fluorescence images of GUV recruitment of ATG13(1-230)-GFP:ATG101 (100 nM) and WIPI3 (200 nM). The membrane recruitment of ATG13/101 was significantly weakened in the absence of the WF finger and CTD motifs of ATG101. The scale bar is 10 µm. **(E)** Quantitation of ATG13(1-230):ATG101 and ATG13(1-230):ATG101-mutants binding on GUV at different concentrations. ****P < 0.0001 and ns: not significant. All data are mean ± s.d.

Many of the proteins involved in autophagy bind preferentially to highly curved membranes(*35–37*). To assess the role of membrane curvature in this recruitment, we performed a similar liposome binding assay as above with GST-immobilized ATG13^HD^:ATG101 in the presence of Atto647-labeled high curved small unilamellar vesicles (SUVs) (Fig. S2). Because their high curvature exposes packing defects that promote membrane insertion by small hydrophobic motifs, SUVs are more permissive for membrane binding, allowing some normally required biological signals to by bypassed(*38*). In the SUV experiments carried out at the superphysiological concentration of 1 μM protein, we observed substantial recruitment of liposomes to wild-type ATG13^HD^:ATG101 even in the absence of WIPI3. The HORMA dimer was recruited non-specifically to SUVs containing charge-equivalent mole fractions of anionic lipids (3% PI3P, 3 % PI4P, or 9% phosphatidylserine (PS) (Fig. S2A and S2C). However, the same experiments performed using the GST alone or membrane-binding deficient ATG101^ΔCTH^ (WF|DD) showed almost no SUV recruitment (Fig. S2A and S2C). These data show that the ATG101 WF finger and CTH have an inherent ability to bind non-specifically to highly curved acidic membranes when binding is driven by superphysiological protein concentrations. These findings are consistent with the absence of a specific PI3P binding site on the ATG13:ATG101 HORMA dimer itself.

### ATG13–ATG101–WIPI3 complexes stably interact with membrane in MD simulations

We examined the stability of the ATG13-ATG101-WIPI3 and ATG13-ATG101-WIPI3-WIPI2 complexes and their membrane binding capacity using atomistic MD simulations. We placed the complexes, as modeled by AlphaFold2, on a phospholipid bilayer. The ATG13–ATG101–WIPI3 complex remained stable throughout 1 μs MD simulations (Fig. 3A and Fig. S3) across all 4 replicas tested. The conserved LRRG motif of WIPI3 formed persistent interactions with a PI3P lipid (Fig. 3A), mediated by strong electrostatic attraction between the two Arg residues and the PI3P head group. The WF finger and the CTH of ATG101 remained stably embedded in the membrane (Fig. 3A), consistent with their importance for membrane recruitment of ATG101. Throughout the simulations, the C-terminal loop (residues 190–230) of ATG13 maintained its interaction with WIPI3 (Fig. 3B and S3A). Asp217 of ATG13 formed dynamic ion pairs with Lys42 and Lys44 of WIPI3 (Fig.3C, 3D, S3B and S3C). The DHF motif (residues 213-215) of ATG13 persistently interacted with a pocket formed by Val35, Lys44, Arg62, Cys63, Asn64, Asp87, and Leu88 of WIPI3. The Root Mean Squared Deviation (RMSD) of these residues from the initial AlphaFold2 structure remained below ∼1 Å over the course of the simulation (Fig. 3E and S3D).

**Figure 3:**
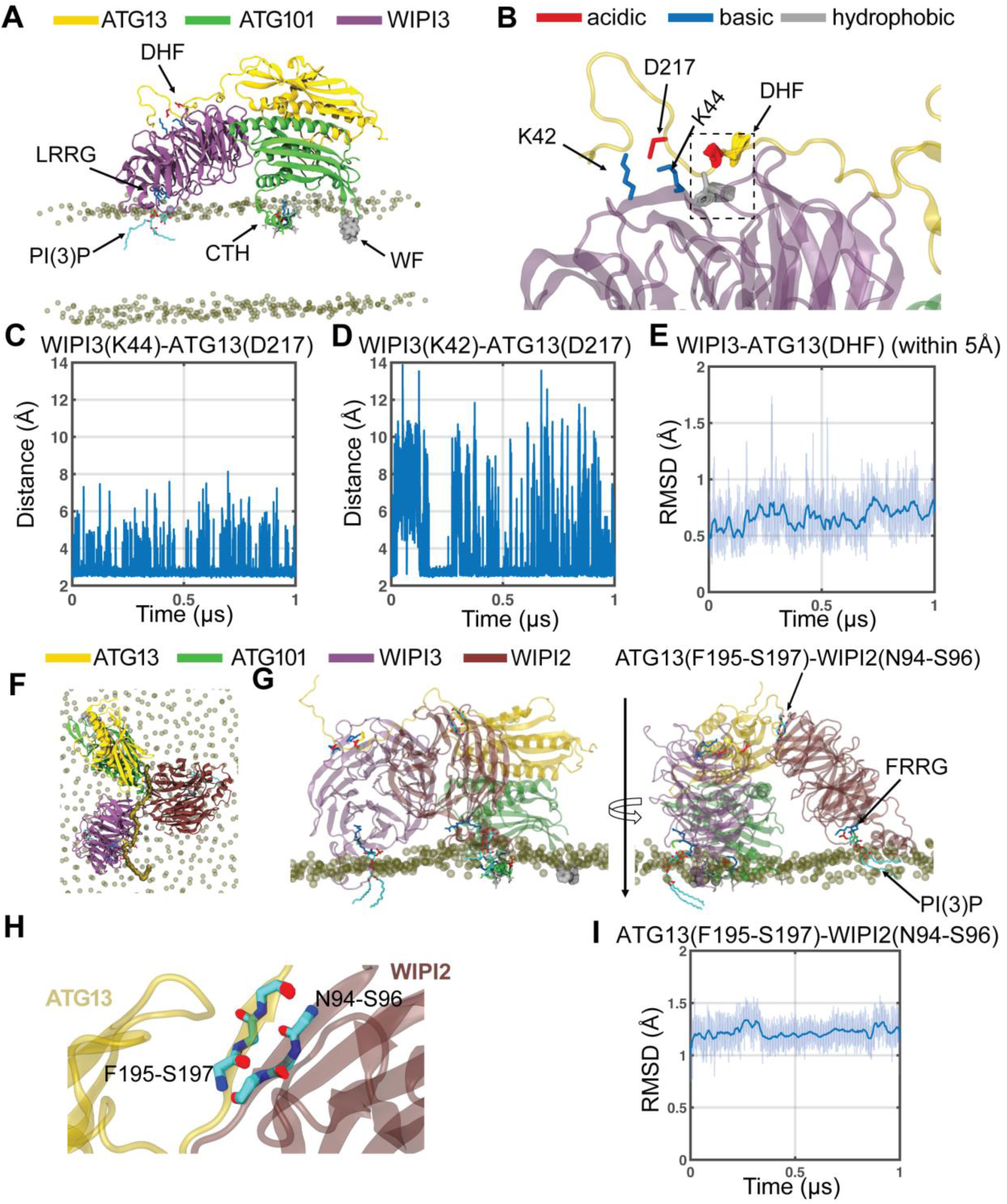
Atomistic MD simulation of membrane-bound ATG13-ATG101-WIPI3 and ATG13-ATG101-WiPI3-WIPI2 complexes. **(A)** Final structure of the ATG13–ATG101–WIPI3 complex after a 1 μs simulation. The ATG101 WF finger (spheres) and C-terminal amphipathic helix (CTH; sticks, color-coded by residue type) are embedded in the membrane, while the WIPI3 LRRG motif interacts with a PI3P lipid (P3 atom shown as an ice-blue sphere). Only phosphate groups of membrane lipids are shown for clarity. **(B)** Zoomed-in view of the ATG13–WIPI3 interface. Dynamics of the DHF motif are illustrated by superimposing conformations sampled at 1 ns interval over the final 500 ns of 1 μs simulation trajectory. Salt bridges between ATG13 (D217) and WIPI3 (K42, K44) are shown as sticks. **(C– D)** Time evolution of the minimum distance from ATG13 D217 (atoms OD1, OD2) to WIPI3 K44 (**C**) and K42 (**D**) NZ atoms. **(E)** Backbone Root Mean Squared Deviation (RMSD) of the ATG13 DHF motif and WIPI3 residues within 5 Å of the DHF. **(F–G)** Top (**F**) and side (**G**) views of the ATG13-ATG101-WIPI3-WIPI2 complex on a membrane after a 1 μs simulation. The ATG13 C-terminal unstructured region mediating interactions with WIPI3 and WIPI2 is in surface representation. The WIPI2 FRRG motif (sticks, color-coded by residue type) interacts with a PI3P lipid (G, right). **(H)** Zoomed-in view of the ATG13–WIPI2 interface, showing backbone atoms of interacting residues (ATG13 F195–S197; WIPI2 N94–S96) sampled every 1 ns during the final 500 ns of the trajectory. **(I)** Backbone RMSD of the residues in (H) as a function of simulation time. Snapshots and plots in panels (B-E) and (H-I) for additional replica simulations are provided in Fig. S3.

To assess the consistency of our AlphaFold prediction model, we performed multiple runs of Alphafod of ATG13-ATG101-WIPI3. In 500 runs (10 diffusion samples each for 50 random seeds) we observed high structural convergence, with the described above interface appearing in 97% of the models (485 out of 500), forming a dominant cluster with a backbone RMSD of ATG13-WIPI3 interface < 1.7 Å (Fig. S4A). In this primary cluster, the ATG13 IDR consistently utilizes its DHF motif to bind WIPI3, stabilized by salt bridges between ATG13 D217 and WIPI3 K42/K44. The high confidence of this interface is further supported by low Predicted Aligned Error (PAE) scores (Fig. S4B). We identified only 15 outlier structures (3%) featuring alternative binding modes. Notably, one outlier class (black dashed box, Fig. S4A) maintains the ATG13-WIPI3 interface but exhibits a misaligned LRRG motif and WF finger. This non-planar orientation would sterically preclude the simultaneous membrane binding of ATG101 and WIPI3, rendering it biologically implausible. To further evaluate the intrinsic stability of these interactions, we performed additional MD simulations of the protein domains in complex with their respective peptide binders in a solvent-only environment. By removing the membrane, we enabled unrestrained binding and enhanced the sampling of the conformational space. A fragment complex (residue 185-284 of WIPI3 and 210-220 of ATG13) derived from an outlier (orange dashed box, Fig. S4C) dissociated within 185 ns (Fig. S4D), confirming that rare AlphaFold-3 predictions correspond to unstable, low-affinity states. By contrast, the fragment complex representing the dominant cluster (residue 18-110 of WIPI3 and 210-220 of ATG13) remained stable throughout a 1 μs simulation (Fig. S4E), maintaining the DHF motif orientation and the D217–K42/K44 salt-bridge network. Finally, MD simulations of the complete ATG13-ATG101-WIPI3 hetero-trimer confirmed a stable association for 1 μs, even in the absence of membrane anchoring (Fig. S4F). The high convergence rate of AlphaFold-3, the low PAE scores, and the MD-verified stability of the dominant cluster—contrasted with the rapid dissociation of outliers— collectively provide a high level of confidence in the proposed ATG13-ATG101-WIPI3 structural model.

We sought to test the impact of ATG101 WF motif on the stability and membrane association of ATG13-ATG101-WIPI3 complex by MD stimulation. Our simulations revealed that the substitution of the hydrophobic WF finger with the negatively charged DD residues leads to a rapid loss of membrane anchoring (Fig. S5A, S5B). The DD segment detaches from the lipid bilayer and becomes fully solvent-exposed within the first 100 ns of simulation. This localized unbinding event imposes significant structural strain on the hetero-trimeric assembly. Specifically, the dissociation of the DD segment triggers a substantial perturbation of the ATG13-WIPI3 interface (Fig. S5C, S5D). The membrane detachment of the DD segment from the membrane and the resulting change in relative protein orientation affected also the ATG13-WIPI3 interaction mediated by the DHF motif. We observed a sharp increase in the RMSD (Fig. S5E) of the interface residues (the DHF motif and its immediate WIPI3 contacts), which rose to 4–5 Å following the loss of the DD–membrane anchor. Notably, while the C-terminal amphipathic helix of ATG101 remains membrane-bound, it is insufficient to maintain the overall orientation or the structural integrity of the ATG13-WIPI3 interface. These simulations demonstrate that the WF finger is not only essential for membrane recruitment and thereby stabilizes the entire hetero-trimeric assembly. The observed partial unbinding and subsequent interfacial disruption provide a clear mechanistic rationale for the loss-of-function observed in experimental mutants.

Further, the ATG13–ATG101–WIPI3–WIPI2 complex also remained stably associated with the membrane surface (Fig. 3F). The FRRG motif of WIPI2 engaged persistently with a PI3P lipid molecule (Fig. 3G). ATG13 formed a stable β-strand interaction with WIPI2 through residues F195–S197 of ATG13 and N94–S96 of WIPI2 (Fig. 3H and S3E). The RMSD of these interacting residues remained ∼1-1.5 Å (Fig. 3I and Fig. S3F), confirming a robust and stable ATG13–WIPI2 interface. All the molecular interactions described above for WIPI3 were preserved (Fig. S3G-S3K) when we extended the system to include simultaneous binding of WIPI2 and WIPI3.

### Reconstitution of the recruitment and activation of ULK1 at membranes

Having established a role for the ATG101 subunit in recruiting the ULK1 complex to membranes, we wondered whether loss of membrane binding by ATG101 would manifest a functional defect in the catalytic activity of ULK1. To assay this, we reconstituted the phosphorylation of membrane-associated ATG16L1 by full-length ULK1 in the context of its complex with ATG13:ATG101, the FIP200 structural core (residues 1-640)(*13*) in the presence of PI3P-containing liposomes, WIPI2, and WIPI3. We used a minimal substrate of the ATG16L1(78-300) fragment that contains the ULK1 phosphorylation site Ser278(*20*) and binding sites for both WIPI2 and WIPI3(*39, 40*)(Fig. 4A). As a readout of ULK1 activity, we used a phospho-specific antibody against ATG16L1 phosphoserine 278(*20*). In the absence of liposomes, the catalytic activity of ULK1 was minimal. Upon addition of liposomes containing 3% PI3P and WIPI2 and WIPI3, we saw a large time-dependent increase in ULK1 phosphorylation of ATG16L1 (Fig. 4B and 4C). Removal of WIPI2 or WIPI3 resulted in a significant decrease in ATG16L1 phosphorylation. Experiments performed using ATG13 constructs containing W3IR motif mutants (ATG13(HF|DD)) and ATG101 membrane-binding deficient constructs (ATG101(WF|DD)) both showed dramatically reduced ULK1 catalytic activity (Fig. 4B and 4C), demonstrating that ULK1 activity against a membrane associated substrate strongly depends on the synergistic recruitment of the complex to the membrane by both ATG13:WIPI binding and membrane docking by the ATG101 WF finger.

**Figure 4.**
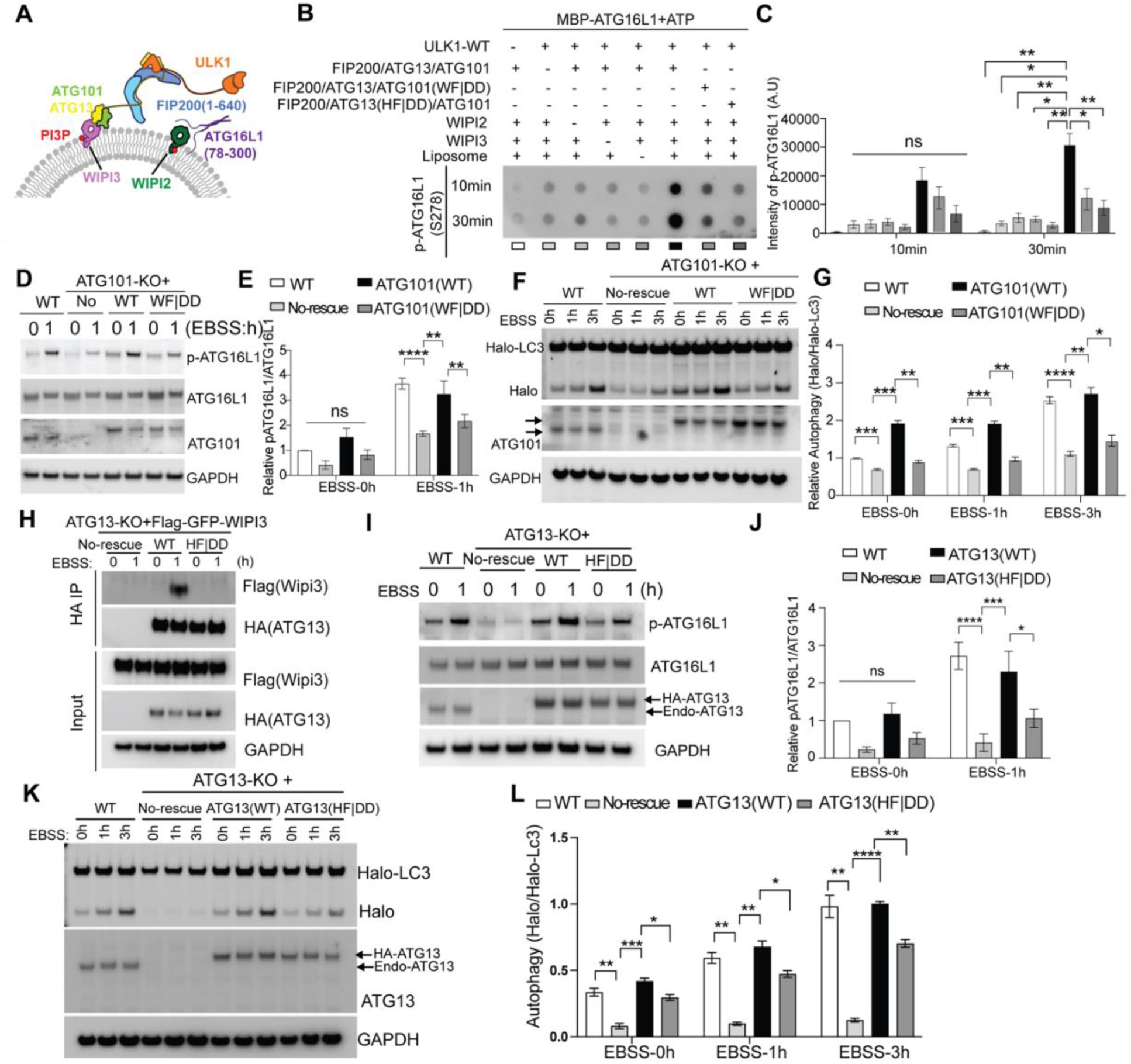
ULK1 complex activity depends on WF finger of ATG101 and W3IR of ATG13. **(A)** Schematic of the kinase assay depicting each of the components. **(B, C)** The dot-blot showed the p-ATG16L1 signal in indicated conditions. And the quantification from 5 independent biological repeats was shown in (C). statistics: Two-way ANOVA + Sidak’s multiple comparisons test**. (D, E)** ATG101 knockout (KO) cells without rescue or rescued with different versions of HA-tagged ATG101 were treated with EBSS for indicated times. P-ATG16L1 signal was analyzed by immunoblotting (IB) (D); quantification of relative p-ATG16L1/ATG16L1 were shown in (E), n= 5 biological replicates. **(F, G)** ATG101 knockout (KO) cells without rescue or rescued with different versions of HA-tagged ATG101 were incubated with 100 nM Halo ligand for 30 min. After this, cells were treated with EBSS for indicated times before harvest and analyzed by immunoblotting (IB) (F), and the relative ratio of the cleaved Halo band was quantified (G), n=5 biological replicates. **(H)** Cells as indicated were transfected with Flag-GFP-WIPI3 for 24h, then treated with or without EBSS and subjected to HA-immunoprecipitation**. (I,J)** ATG13 knockout (KO) cells without rescue or rescued with different versions of HA-tagged ATG13 were treated with EBSS for indicated times. P-ATG16L1 signal was analyzed by immunoblotting (IB) and the quantifications were shown in (J). n=5 biological replicates. **(K, L)** ATG13 knockout (KO) cells without rescue or rescued with different versions of HA-tagged ATG13 were incubated with 100 nM Halo ligand for 30 min. After this, cells were treated with EBSS for indicated times before harvest and analyzed by immunoblotting (IB) (K), and the percentage of the cleaved Halo band was quantified (L), n=4 biological replicates. Statistics for (E), (G), (J) and (L): Two-way ANOVA + Tukey’s multiple comparisons test. Mean ± SEM was shown. *P<0.05, **P<0.01, ***P<0.001, ****P<0.0001, ns, not significant.

We sought to confirm our *in vitro* results in a cellular model. We first investigated the role ATG101 WF finger in autophagy. Consistent with the *in vitro* results, we found complementation of ATG101-WT but not ATG101(WF|DD) rescue the loss of p-ATG16L1 in ATG101-KO background (Fig. 4D and 4E). Rescue with ATG101(WF|DD) in ATG101-KO cells exhibited a significant loss of autophagic flux compared to ATG101-WT (Fig. 4F and 4G), which is consistent with previous work (*12*). In probing the ATG13:WIPI3 interaction in cells, we found that starvation induces a detectable interaction of ATG13-WT, but not ATG13(HF|DD), with WIPI3 (Fig. 4H). As in the reconstituted biochemical assay, we found the ATG13(HF|DD) mutant displayed lower cellular kinase activity, as monitored by pSer278 ATG16L1 levels in cell lysates (Fig. 4I and 4J). Next, we characterized the effect on autophagy via a Halo-LC3 flux assay(*41*). As shown in Fig. 4K and 4L, complementation of ATG13(HF|DD) in ATG13-KO cells exhibited a significant loss of autophagic flux compared to ATG13-WT highlighting the importance of ATG13-WIPI3 interaction for autophagy.

The ULK1 complex is not only a master regulator of general autophagy but is also required for iron-chelation-induced mitophagy (*42*). Therefore, we sought to determine whether the ATG13(HF|DD) mutant affects this pathway. To monitor mitophagy we used the well-characterized *mito*-QC assay in ARPE-19 cells, which was previously found to undergo robust mitophagy (*42, 43*). This assay uses cells that express a tandem mCherry–GFP tag fused to the outer mitochondrial membrane localization signal derived from FIS1 (residues 101–152). As GFP signal is quenched under acidic lysosomal conditions, mitochondria that have been delivered to lysosomes (referred to as mitolysosomes) can be identified as red-only puncta and quantified. *mito*-QC ARPE19 cells, overexpressing ATG13 WT or the ATG13(HF|DD) mutant, were treated with the iron chelator deferiprone (DFP), which mimics hypoxia and induces HIF1-dependent transcription of the mitophagy receptors BNIP3 and NIX (also known as BNIP3L) and ULK1-dependent mitophagy (*43, 44*). Overexpression of ATG13(HF|DD) significantly impaired mitophagy compared to cells overexpressing ATG13 WT (Fig. S6A, S6B), consistent with the results shown for general autophagy in the ATG13 KO rescue cells (Fig. 4K, 4L). Overexpression of ATG13 did not markedly alter the levels of other ULK1 complex components (Fig. S6C). As expected, DFP treatment in WT cells stabilized HIF1α and reduced the levels of mitochondrial proteins such as OMI, a serine protease localized to the mitochondrial intermembrane space that can be used as a marker of mitochondrial content(*42, 43*), in control cells, consistent with the induction of mitophagy (Fig. S6C). Although HIF1α was stabilized in HF|DD mutant cells, OMI was reduced in a less extent than WT cells. These results suggest that the ATG13(HF|DD) mutant acts in a dominant-negative manner. Taken together, these results indicated that the interaction between ATG13 and WIPI3 plays an important role in autophagy, including mitophagy.

### ULK1 IDR binds directly to the ATG13 HORMA domain

ULK1 is comprised of three distinct domains, including the kinase domain (KD) from residues 1-290, a ∼500 residue intrinsically disordered region (IDR), and a C-terminal tandem MIT/EAT domain that forms a folded complex with the N-terminus of FIP200 and the C-terminal MIM domain of ATG13 (Fig. 5A)(*13*). Even when the ATG13:ATG101 HORMA dimer is tightly engaged with membranes, the ULK1 KD is tethered only distantly from the membrane by the ULK1 IDR and a further > 150 residues of ATG13 IDR. We speculated that a mechanism must exist to bring the ULK1 kinase closer to the membrane to reach its membrane-bound substrates such as ATG16L1.

**Figure 5.**
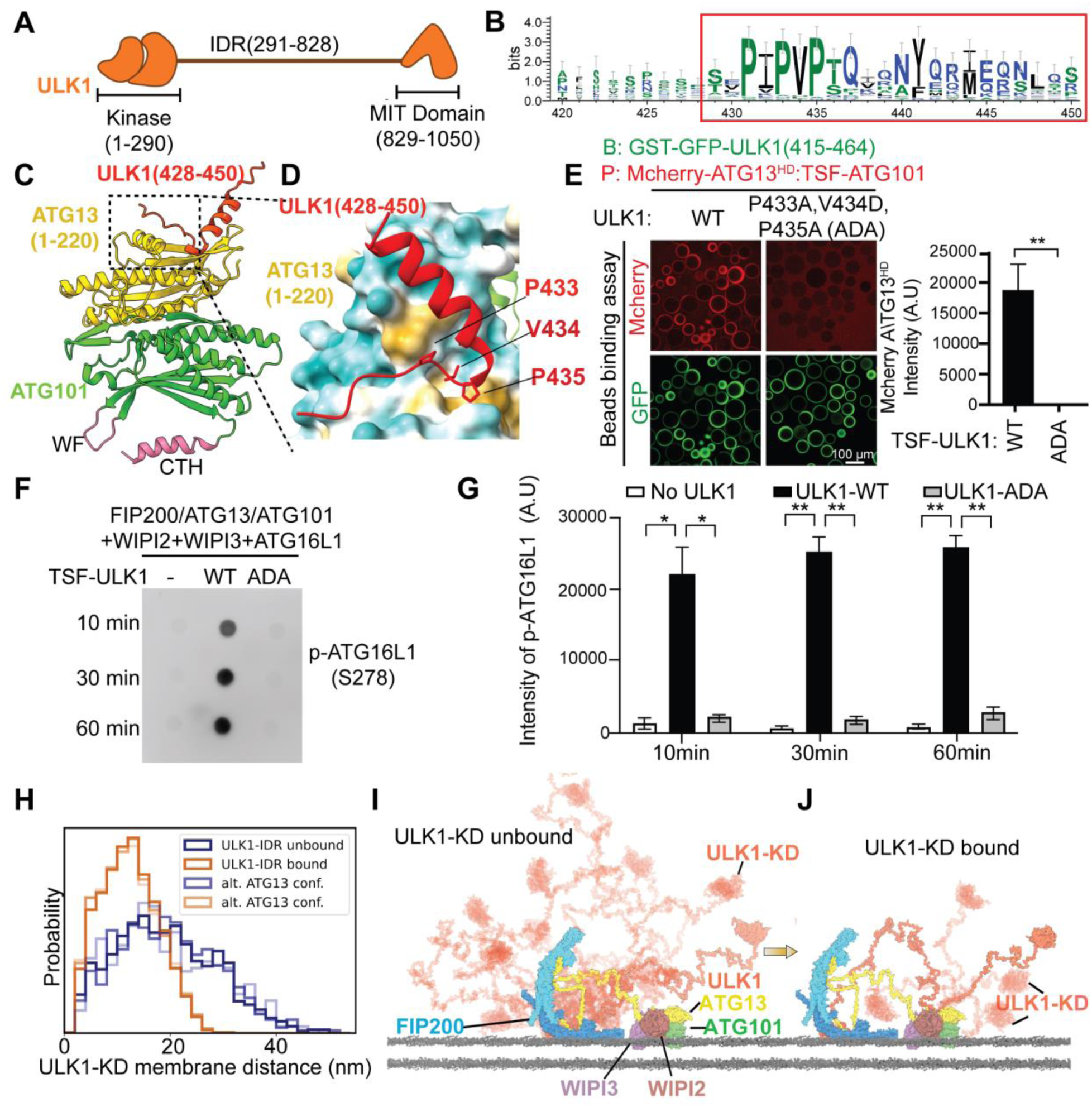
ULK1 binds to ATG13HD/ATG101 through its IDR and this interaction is important for kinase activity. **(A)** ULK1 schematic. **(B)** Sequence logo displaying conservation patterns within the IDR. The sequences in red box indicated the potential ATG13 binding motif. **(C)** AlphaFold2 structural model of ATG13(1-220)-ATG101 and ULK1 peptides. **(D)** Close up view of the interface between the residues P433, V434 and P435 of the ULK1 with ATG13. **(E)** GST-GFP-ULK1(415-464) constructs that contain triple mutation (P433A, V434D, and P435A, ADA) in ULK1 fail to recruit mCherry-ATG13^HD^:TSF-ATG101. The intensity of mcherry channel was quantified and shown as bar graph. n=5 biological replicates, statistics: Unpaired t-test. **(F, G)** Dot-blot and quantification showed that full length ULK1 containing the ADA mutant almost completely lost its catalytic activity. “B” indicated bait, and “P” indicated prey. n=4 biological replicates, statistics: Two-way ANOVA + Sidak’s multiple comparisons test. **(H)** Histograms of ULK1-KD distance to the membrane in presence and absence of ULK1-IDR binding to ATG13. The distributions were obtained from ULK1-IDR ensembles with three alternative conformations of ATG13. **(I)** 10 conformations of ULK1C including FIP200 dimer, full-length ATG13, full-length ULK1, WIPI2-3 and ATG101 in absence of ULK1-IDR interaction with ATG13. One conformation is shown as non-transparent, the other 9 as transparent. **(J)** 5 conformations of ULK1C in presence of ULK1-IDR interaction with ATG13. Mean ± SEM was shown. *P<0.05, **P<0.01, ***P<0.001.

To probe whether there were any additional interactions between the ATG13:ATG101 dimer and ULK1, we again used AlphaFold2 to predict interactions between them. We predicted a direct, evolutionarily conserved interaction of the ULK1 intrinsically disordered region (IDR) (Fig.5B-5D, S7A and S7B) with ATG13^HD^. The interaction is predicted to be driven predominantly by a stretch of conserved proline residues and hydrophobic amino acids encompassing residues 428-450 of ULK1 (Fig.5C, 5D). Again, to assess the consistency of our AlphaFold prediction model, we performed multiple runs of Alphafold of ATG13-ATG101-ULK1 (428-450). Out of 500 runs, outliers were observed only twice, indicating a very high degree of consistency in the prediction (Fig. S7C-S7E). The binding site is in a cleft that is formed by the β-strand just N-terminal to the ATG13 CTH and the second alpha helix of ATG13 (Fig. 5D). To confirm the existence of this direct interaction between ATG13^HD^ and ULK1, we performed a GST-resin based interaction assay using GST-ATG13^HD^:ATG101, and mCherry labeled ULK1 constructs lacking the ULK1 EAT domain, including mCherry-ULK1^ΔEAT^ (1-828), mCherry-ULK1 ^ΔΚDΔEAT^ (291-828), and ULK1^KD^ (1-290) (Fig.S8). Consistent with the AlphaFold2 prediction, the mCherry-ULK1 construct mCherry-ULK1 ^ΔΚDΔEAT^ containing only the ULK1 IDR bound to the HORMA dimer (Fig.S8A and S8B), while ULK1^KD^ did not (Fig. S8C). We selected three of the highly conserved residues at the interface between ATG13 and ULK1, P433, V434, and P435 (Fig. 5B and 5D), for further functional analysis using site-directed mutagenesis. Using a GST-tagged fragment of the ULK1 IDR encompassing residues 415-464, and a construct containing mCherry-ATG13^HD^:ATG101, we found that constructs containing the triple mutation of ULK1 P433A/V434D/P435A (hereafter, ADA) showed a complete loss of binding (Fig. 5E).

To test the role of the ATG13^HD^ -ULK1 interaction in ULK1 complex catalytic activity, we again used an anti-pSer278 ATG16L1 phosphospecific antibody as a probe (Fig. 4A). We observed that the ULK1 ADA mutation almost completely lost the ability to phosphorylate ATG16L1 at this site (Fig. 5F and 5G). We conclude that the consequence of the ADA mutation is to severely impair nanoscale ULK1 kinase co-localization with its substrates.

### The ULK1-IDR interaction with ATG13^HD^ drives proximity of ULK1 KD to the membrane

To model the effect of the ULK1 intrinsically disordered region (IDR) mutants we generated conformational ensembles of ULK1C in presence and absence of the interaction between ULK1 residues 433-435 and the ATG13 HORMA domain. The AlphaFold2 model of the interaction shows V434 and P435 inserted into a hydrophobic pocket on ATG13 (Fig. 5D). For the rest of the complex, we generated a model of the ULK1C core including IDR regions and the ULK1 KD as assembled in 2:1:1 stoichiometry consistent with the experimental structure (*13*) (Fig. S9). We modeled the ATG13-IDR such that ULK1 C927 and C1003 were engaged with the membrane (see Methods for details) as has been recently reported (*45*). Average distances between the ULK1-KD and the membrane show a reduction (19 nm to 12 nm) upon binding of the IDR to ATG13, an effect that is consistent across multiple conformations of ATG13 (Fig. 5H). This is illustrated by the reduced conformational space accessible to the complex (Fig. 5I, 5J). The loss of ATG13^HD^ and ULK1 interaction upon mutation thus increases the average distance to the membrane and thereby to the membrane-bound substrates of ULK1.

### PVP mutation of ULK1 affects mitophagy and autophagy

To confirm the importance of the ATG13 HORMA-ULK1 interaction in a cellular setting, we examined starvation-induced autophagy and iron-chelation-induced mitophagy. To monitor starvation-induced autophagy we used ULK1/2 double knockout MEFs (*46, 47*) stably expressing the tandem mCherry-GFP-LC3 autophagy reporter, which works along a similar principle to the previously mentioned *mito*-QC assay (Fig. S10). Into these cells, we re-introduced FLAG-tagged wild-type (WT) ULK1 or the ULK1-ADA mutant via retroviral-mediated transduction and induced autophagy by incubating cells in amino acid-free Earle’s Balanced Salt Solution (EBSS). As can be seen, reintroduction of ULK1 restored autophagic flux in the ULK1/2 DKO cells (Fig. S10A, S10B). However, the level of autophagy induced by the ULK1 ADA mutation was significantly less compared to WT ULK1, implying ULK1’s interaction with the ATG13 HORMA domain is critical for efficient autophagy. In parallel, the ULK1 ADA mutant displayed lower cellular kinase activity, as monitored by pSer278 ATG16L1 levels in cell lysates (Fig. S10C, S10D).

We next evaluated the effect of ADA mutation on mitophagy. Using ULK1-KO ARPE-19 *mito*-QC cells (*44*), we stably reintroduced FLAG-tagged WT ULK1 or the ULK1-ADA mutant and induce mitophagy with DFP. While the levels of ULK1 re-expression were higher than endogenous levels seen in WT cells, both the FLAG-tagged WT and ADA mutant were at comparable levels and did not alter the abundance of other ULK1 complex members (Fig. S10E). Re-expression of ULK1-WT caused a robust increase of the number of mitophagic cells upon DFP treatment, which was significantly higher compared to cells re-expressing the ULK1-ADA mutant (Fig. 6A, B). To monitor mitophagy independently of the *mito*-QC reporter, we also examined mitochondrial protein levels by western blot (Fig 6C, 6D and S10E). The effect of DFP on stabilizing HIF1α was confirmed and, consistent with the *mito*-QC assay, the level of mitochondrial OMI protein was significantly reduced following DFP treatment in ULK1-WT expressing cells but not in cells expressing either the empty vector (ULK1-KO) or the ULK1-ADA mutant. This shows that not only is ULK1 essential for DFP-induced mitophagy, but its interaction with the ATG13 HORMA domain plays a critical role.

**Figure 6:**
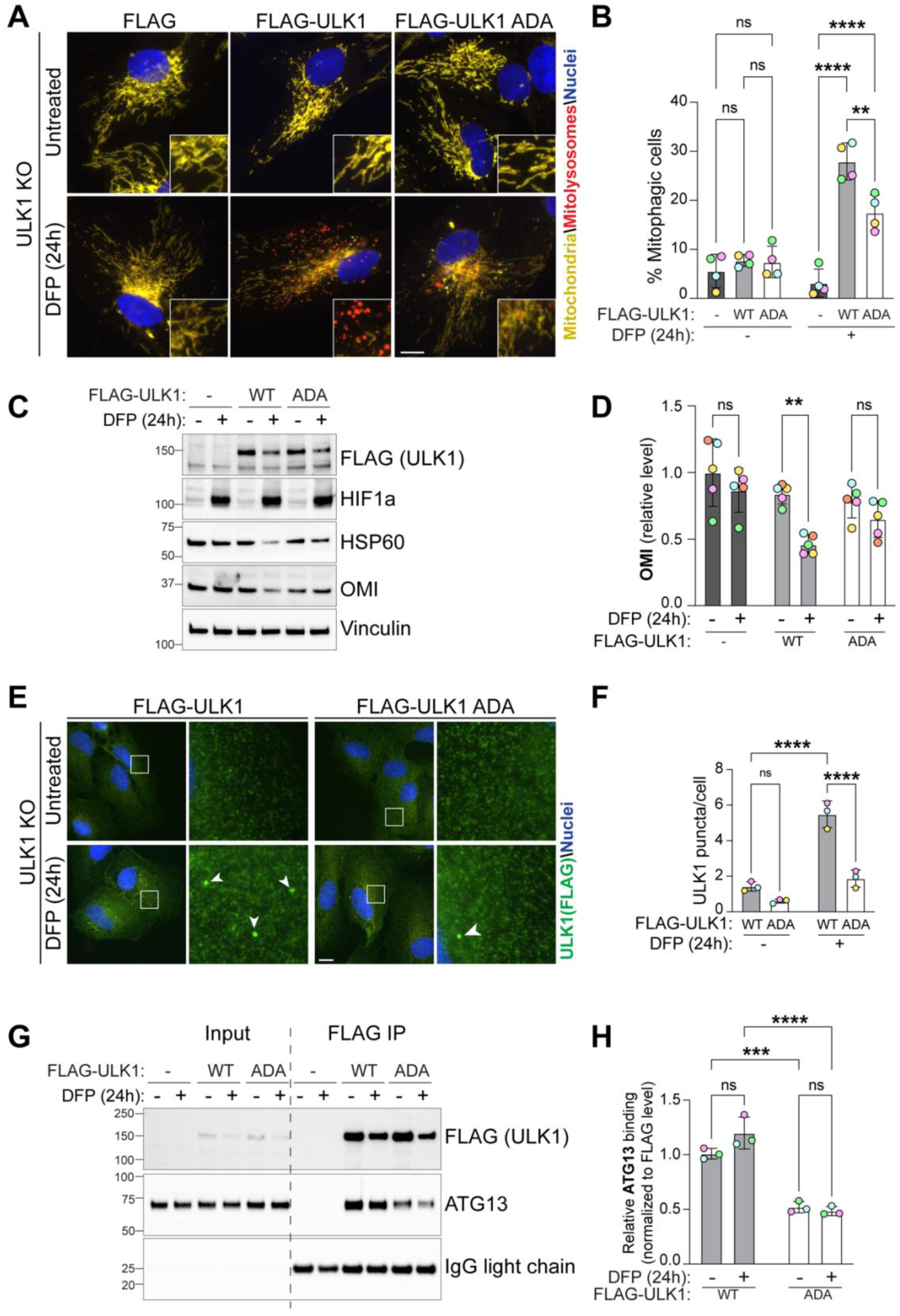
PVP motif of ULK1 is essential for mitophagy. (**A**) Representative wide-field images of ULK1 knock-out (KO) ARPE-19 cells stably expressing an empty vector (FLAG), WT or ADA mutant FLAG tagged ULK1, and the *mito*-QC reporter. Cells were treated with 1mM DFP for 24 hours prior to imaging. (**B**) Flow cytometry analysis of the mCherry/GFP ratio (n = 4 biological replicates); statistics: two-way ANOVA with Tukey’s multiple-comparisons test. **(C, D)** Representative immunoblots and quantification of the indicated proteins from cells as in (A) (n = 5 biological replicates); statistics: two-way ANOVA with Tukey’s multiple-comparisons test. (**E**) Representative wide-field images of cells as in (A), treated with 1mM DFP for 24 hours and immunostained with anti-FLAG (green). (**F**) Quantification of the puncta per cells from 3 independent experiments with a minimum of 70 cells being analyzed per condition in each experimental replicate; statistics: Two-way ANOVA with Tukey’s multiple comparisons test. (**G, H**) Cells as in (A), were treated with 1mM DFP for 24 hours and subjected to FLAG-immunoprecipitation. Representative immunoblots (G) and quantification of the indicated proteins (H) are shown (n=3 biological replicates); statistics: Two-way ANOVA with Tukey’s multiple comparisons test. Data information: Enlarged images are shown in the lower corners. Nuclei were stained in blue (Hoechst) and scale bars: 10 µm. All data are mean ± s.d.; Statistical significance is displayed as **P ≤ 0.01; ***P ≤ 0.001; ****P ≤ 0.0001; ns, not significant.

To examine the cellular context of the ATG13 HORMA-ULK1 interaction in more detail, we analyzed ULK1 puncta formation upon mitophagy induction. ULK1 localizes to sites of autophagosome formation upon mitophagy induction, which can be monitored by immunofluorescence microscopy (Fig.6E, 6F). DFP strongly induced WT ULK1 puncta formation; however, this was significantly impaired in the ULK1-ADA mutant expressing cells. To support these data, we also immunoprecipitated FLAG-tagged ULK1 from our cells under mitophagy-inducing conditions and examined the co-IP of ATG13 (Fig 6G, 6H). We found that co-IP of ATG13 with the ULK ADA mutant was reduced compared to that with WT ULK1. These data show that the interaction of ULK1 with the ATG13 HORMA domain is important for autophagy and mitophagy because of its role in the recruitment and stabilization of the ULK1C at initiation sites.

## Discussion

The recruitment to, and activation of, ULK1 on membranes is a central event in autophagy initiation. Yet how ULK1 recruitment is triggered and how it is coordinated with the activity of the other key initiation complex, PI3KC3-C1, has not been explained. The mechanistic basis for the more than decade-old observation that ULK1C is stabilized at phagophore initiation sites by PI3P(*26*) has been unknown. The strong dependence of autophagy on ATG101(*7*) has not been satisfactorily explained. A major result of the work presented here is to establish a unifying model (Fig. 7) to account for these observations in the autophagy field.

**Figure 7.**
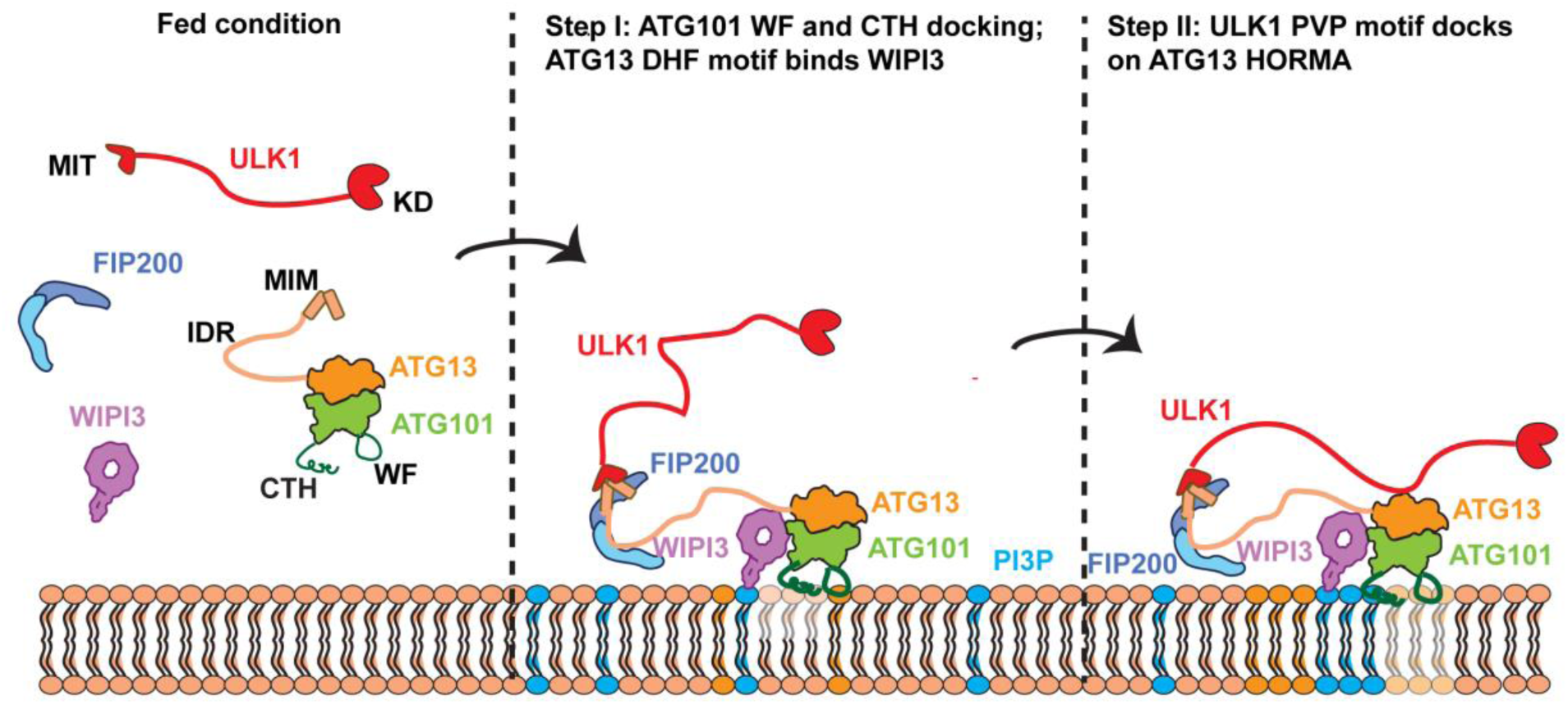
Multistep ATG13:ATG101 and WIPI3-dependent recruitment of ULK1 to membranes in autophagy. When autophagy is initiated, FIP200, ULK1 and ATG13:ATG101 co-assemble. ATG13:ATG101 with the PI3P-binding protein WIPI3 via the ATG13 DHF motif and aligns with the membrane to insert its Trp-Phe (WF) finger into, and CTH onto, the membrane. Next, the ULK1 kinase domain (KD)-proximal PVP motif within the ULK1 IDR docks onto the surface of the ATG13:ATG101 HORMA dimer (HD) and brings the ULK1-KD close to the membrane surface, where its autophagy-related substrates are also located.

It is increasingly clear that autophagy initiation proceeds not in a strict hierarchy, but rather through a series of positive feedback loops in which downstream effects reinforce upstream processes. Examples include PI3P product activation of PI3KC3-C1(*48*) and ATG8 binding to AIM/LIR motifs within autophagy core complexes(*49*). PI3P-dependent stabilization was originally attributed to basic residues within ATG13(*26*), a finding that was first called into question by the absence of a structural PI3P binding site in ATG13(*11, 12*). Here, we showed that ATG13:ATG101 by itself does not bind to PI3P using GUV binding assays. The rigorous reanalysis carried out here, incorporating structural insights, revealed that the mechanism is much more complex and nuanced than originally thought. The synergism between WIPI2, WIPI3, and the membrane interacting WF and CTH motifs of ATG101 offers a rich set of drivers, each of which is needed individually and each of which is potentially subject to regulation in its own right.

These data help explain why ATG101 has a fundamental and required role in autophagy(*7*). One of the few clearly established roles for ATG101 is to participate, together with ATG13, in bridging the ULK1C to ATG9 via binding to the C-terminus of the soluble IDR of ATG9(*15, 50, 51*). Mutation of the ATG9 binding site only partially reduces autophagic flux, however, indicating that binding to the ATG9 C-terminal peptide cannot be its only function. Moreover, the ATG101 WF has been shown to be functionally essential(*12*), yet the WF residues are not involved in ATG9 IDR binding. The identity of the ATG101 WF binding partner has been unresolved for the past decade. Here, we show that the ATG101 WF finger is essential for synergistic membrane recruitment of ULK1C by WIPIs, which seems sufficient to account for its essential role.

Both ATG13 and ATG101 belong to the HORMA domain protein family(*52*), and other members of this family have been shown by x-ray crystallography to undergo a slow metamorphic refolding that alters their interaction capacity. Despite a hypothetical proposal that ATG13 and ATG101 can undergo metamorphosis (*51*), a putative event that takes place on a time scale orders of magnitude slower than autophagy itself, there has been no experimental structural validation for this idea. The structural models and findings presented here correspond to the established crystal structure of human ATG13:ATG101(*11*). None of the findings described here depend on conformational metamorphosis.

The findings described here add another role to the expanding repertoire of WIPI functions in autophagy (*53*). WIPIs are central to the regulation of ATG8ylation by virtue of the role of WIPI2, and to some extent, WIPI3 in the recruitment of the ATG12-ATG5-ATG16L1 complex by virtue of their binding to a helical motif as well as additional sites in ATG16L1(*31, 39, 40*). WIPIs, in particular WIPI4, play a central role in autophagic membrane expansion by promoting ER-phagophore membrane contact sites and stabilizing bridge-like lipid transport from the ER to the phagophore by ATG2(*54–56*). WIPI3 has been less intensively studied than WIPI2 and WIPI4 (*53*), despite that human genetic evidence implicates nonsense mutations of WIPI3 in intellectual disability (*57–59*), WIPI3 was found to co-IP and to colocalize with ULK1C subunit FIP200 upon induction of autophagy by starvation (*60*), consistent with our findings. WIPI3 is important for starvation-induced autophagosome formation in melanoma cells, and essential for etoposide-induced alternative autophagy in *Atg5^KO^*MEFs, respectively(*59, 60*). In contrast to most other work in the field, WIPI3 knockout by CRISPR was found to have no effect on starvation-induced autophagy flux in HEK293 cells (*61*). The reason for this difference is unclear and will require further study. Apart from one report (*61*), our own findings and the preponderance of the published data are consistent with a role for WIPI3 in human health, in ULK1C recruitment to sites of autophagy initiation, and in multiple forms of autophagy.

The observation of a membrane-anchoring role for the ATG13:ATG101 HD dimer leads naturally to the question of positioning of the catalytic KD of ULK1 itself. On the nanoscale, the combined ∼650 residues of IDR contributed by ATG13 and ULK1 leave the KD free to explore a volume of ∼6x10^4^ nm^3^, estimated based on a maximum membrane-KD separation of 40 nm. The maximum separation is cut in half to ∼ 20 nm by the HD-ULK1 IDR interaction, reducing the volume available to explore by ∼8-fold. This is equivalent to a one order of magnitude increase in the local concentration of the ULK1 KD in the vicinity of the membrane. This implies that simultaneously, the ULK1 KD must be removed from the layer 20-40 nm away from the membrane, which might prevent inappropriate phosphorylations of proteins farther from the phagophore. This distance scale corresponds to a typical membrane contact site. The ULK1 IDR-HD interaction might serve to confine the ULK1 KD to the phagophore surface and prevent undesired reactions on the cognate surface of the ER. Clusters of up to 160 ULK1 molecules(*24*) have been implicated in autophagy initiation, but the mechanism of clustering has not been defined. The ULK1 IDR-HD interaction could also serve as a mechanism to cluster multiple ULK1C complexes in *trans* at a phagophore initiation site. Our observation that the ULK1 ADA mutant significantly reduces the number of ULK1C puncta observed, at the light microscopy level, upon mitophagy induction supports this hypothesis.

While the subunits of ULK1C lack any of the common conserved membrane-interacting motifs such as FYVE, PH, and C2 domains, there are other documented membrane interactions. The ULK1 EAT domain contains two conserved surface-exposed Cys residues that are subject to palmitoylation upon autophagy induction(*45*). At least in the case of the 2:1:1 stoichiometry ULK1C core assembly, we found that ULK1 EAT Cys palmitoylation is compatible with the membrane recruitment pathway described here. ULK1C can form a physical supercomplex with PI3KC3-C1(*13*). RAB1A binding appears to be the dominant driver of PI3KC3-C1 localization to phagophore initiation sites(*62*). Hypothetically, RAB1A could indirectly contribute to ULK1C recruitment with PI3KC3-C1 acting as bridge. ATG8 proteins are well established to function in scaffolding ULK1 localization, which presumably occurs downstream of ATG16L1 recruitment and activation by ULK1(*63*), and therefore downstream of the events described in our study. The interplay between these various mechanisms will require further investigation.

A final trending theme highlighted by this study is the multistep nature of peripheral membrane protein recruitment and activation (Fig.7). For example, PI3KC3-C1 is initially recruited to membranes in an inactive conformation by RAB1A(*62*), and subsequently activated and stabilized on membranes by the unmasking of a covalent N-myristoyl modification of the N-terminus of its VPS15 subunits(*16*). ATG16L1 is initially recruited by binding of WIPI2 to a region separated from its catalytic domain by a 150-residue coiled coil, which leaves the catalytic unit still remote from the membrane surface in a nanoscale sense(*37*). A set of additional interactions drive the progressive stepwise positioning of the ATG12-ATG5 to promote ATG3-dependent ATG8ylation of membrane-localized phosphatidylethanolamine (PE). mTORC1 is initially localized to the lysosomal membrane by active Rag GTPase dimers, but complete membrane docking and full activation requires additional steps mediated by the Rheb GTPase and membrane-anchoring residues within mTORC1 itself(*64*). ULK1 is analogous to mTORC1 in the sense that both are kinase master regulators of many membrane-associated processes. Thus, it seems fitting that both are recruited and activated through a complex and stepwise process, providing for precise control and regulation at multiple steps.

## Materials and Methods

### Cell culture

HEK293-Lenti-X, Stable HeLa cell lines were maintained in Dulbecco’s modified Eagle’s medium (DMEM, Gibco) supplemented with 10% (v/v) heat-inactivated fetal bovine serum (FBS, Gibco) and 1x anti-anti (Gibco) (Table S1). HEK293F GnTi cells were maintained in FreeStyle™ 293 Expression Medium (Gibco) supplemented with 1% FBS and 1x anti-anti. ARPE-19 cells (ATCC, CRL-2302) were maintained in 1:1 DMEM:F-12 media (Thermo Scientific) supplemented with 10% (v/v) FBS, 2 mM L-glutamine, 100 U/ml penicillin and 0.1 mg/ml streptomycin. ULK WT and ULK1/2 double knockout MEFs (a kind gift from Prof. Craig Thompson, Memorial Sloan-Kettering Cancer Center) (*47*) and 293FT cells were cultured in DMEM (Sigma-aldrich) supplemented with 10% (v/v) FBS, 2 mM L-glutamine, 100 U/ml penicillin and 0.1 mg/ml streptomycin. All cell lines were confirmed mycoplasma negative using MycoAlert Detection kit (Lonza, LT07-318). Cells were treated as indicated conditions and maintained at 37℃ in 5% CO_2_.

### Stable cell lines

ULK1 KO ARPE-19 cells were generated using CRISPR-Cas9 technology. Targeting guides were transfected in cells using 6 ml of GeneJuice (Sigma #70967) per mg of DNA. After 24 h of transfection, cells were selected using 2 mg/ml of puromycin for 24 h and then allowed to recover. Cells were serially diluted to generate single clones and screened using immunoblotting and DNA sequencing. The following gRNA sequence was used for generating knockout clones: ULK1 KO ARPE-19 clones: paired gRNA, sense (GTTCTCCCGCAAGGACCTGAT) in pBabeD_P_U6 vector and anti-sense (GCCACGGTCTCTGTGCCGCCG) in pX335 vector targeting N-terminal. Reporter cells for mitophagy (*mito*-QC) or autophagy (mCherry-GFP-LC3), and cells overexpressing WT ATG13, HF|DD ATG13, WT ULK1 or ADA ULK1, were generated by retroviral infections. To produce retroviral particles, 293FT cells at 60–70% of confluency, were cotransfected directly in the growth media with X-tremeGENE 9 DNA Transfection Reagent (Roche) ratio 3:1, the cDNA of interest, GAG/POL and VSVG vectors (Clontech). Virus was harvested 48 h post-transfection and applied to cells in the presence of 10 μg/ml polybrene and 20 mM HEPES. Cells were selected with 500 μg/ml hygromycin (Source Bioscience) or 2 μg/ml puromycin (Sigma) after 48 h post transduction. Stable pools were used for experiments.

ATG101-KO, ATG101-KO+Halo-LC3B, and ATG101-KO+Halo-LC3B+HA-ATG101-WT or WF|DD HeLa cell lines were established via lentiviral transduction. Briefly, the sgRNA targeting human ATG101 (sgATG101-#1: TGCGCAACTCTGGTGGCGAT; sgATG101-#2: ACCAGAAGAAGAAGTCTCGC) were cloned into the lentiCRISPRv2 vector (Addgene, 52961) for lentivirus production. To generate lentivirus, Lenti-X 293T cells at 60–70% confluency were co-transfected with Lipofectamine 2000 and a plasmid mixture consisting of pLenti-sgATG101#1 or #2, pCMV-dR8.2 dvpr (Addgene #8455), and pMD2.G (VSVG; Addgene #35616) at a mass ratio of 5:3:2. Viral-#1 as well as virual-#2 supernatants were collected 48 h post-transfection and applied together to target cells supplemented with polybrene (10 μg/ml). Following 48 h of transduction, stably integrated cells were selected with puromycin for 3 days, after which surviving cells were pooled and used for downstream experiments.

To generate the Halo-LC3B-expressing line, lentivirus encoding Halo-LC3B was produced from pCDH1-CMV-HT-LC3-SV40-Hygro (Addgene #182045) using the same packaging protocol described above. ATG101-KO cells were transduced with this virus and subsequently selected with hygromycin (500 ng/ml) for 1 week to establish a stable ATG101-KO+Halo-LC3B cell line.

Finally, lentivirus encoding HA-ATG101-WT or the WF|DD mutant was produced and used to transduce the ATG101-KO+Halo-LC3B cells following the same protocol. For these complementation experiments, puromycin selection was omitted; instead, transduced cells were used directly for experiments following confirmation of HA-ATG101 expression by immunoblotting.

HeLa ATG13 KO+HaloTag-LC3 was a gift from Michael Lazarou’s lab. To generate HeLa ATG13-KO+Halo-LC3+HA-ATG13-WT and HeLa ATG13-KO+Halo-LC3+HA-ATG13-HF|DD stable cell lines, retrovirus was used. To produce retroviral particles, 293FT cells at 60– 70% of confluency, were cotransfected directly in the growth media with polyethylenimine/PEI ratio 5: 3:1, the cDNA of interest (pBMN-HA-ATG13-WT and pBMN-HA-ATG13-HF|DD), GAG/POL (addgene: 14887) and pCMV-VSV-G vectors (Addgene: 8454). Virus was harvested 48 h post-transfection and applied to cells in the presence of 10 μg/ml polybrene. After infected with retrovirus for 48h, part of cells was harvested and applied to western blot to check the expression of the target proteins. Stable pools were used for experiments. All the cell lines descripted above were listed in Table S1.

### Plasmids construction

pCAG-GST-ATG13 (6-196) (Addgene, 203559), pCAG-GST-TEVs-WIPI3 (Addgene, 223800), pCAG-GST-TEVs-WIPI4 (Addgene, 223801), pCAG-FIP200(1-640)-TSF (Addgene, 203545), pCAG-ATG101 (Addgene, 189590) and pCAG-TSF-ATG101 (Addgene, 210858) were from the other members of Hurley lab. pBMN-HA-ATG13 (Addgene, 186223), GAG/POL (Addgene: 14887) and pCMV-VSV-G vectors (Addgene: 8454) were from Michael Lazarou’s lab. TSF-GFP-WIPI3, GFP-ATG13 (1–230) (no tag), TSF-ATG101 ΔCTH (1-198), TSF-ATG101ΔCTH (WF|DD), GST-ATG101(WF|DD) and TSF-ULK1 were synthesized via Integrated DNA Technologies. subcloned into the pCAG vector using Cla I/Xho I sites. LentiCRISPRV2 (Addgene, 52961), pCMV-dR8.2 dvpr (Addgene, 8455). MBP-mCherry-ULK1^ΔEAT^ (1-828), MBP-mCherry-ULK1 ^ΔΚDΔEAT^ (291-828), and MBP-ULK1^KD^ (1-290) were subclone into pCAG vector via homologous recombination. The fragment of HA-ATG101-WT or WF|DD was PCR from pCAG constructed mentioned above. The primer used for PCR were: pCDH1-HA-ATG101-F: ATGTACCCATACGATGTTCCAGATTACGCTTCCGGACTCAGATCTCGAGCTCAAGCT TCG ATGAATTGCCGGTCAGAGGTC; pCDH1-HA-ATG101-FL-R: CTACGTACCACCACACTGGGATCCTTAGAGTGCCAGGGTGTCTTTGAT. Then, the PCR produced were subclone into linearized pCDH1-CMV-HT-LC3-SV40-Hygro (addgene: 182045) (XbaI and BamHI) construct via homologous recombination. Q5 site-directed mutagenesis kit was used for generating all the mutant constructs, including pBMN-HA-ATG13-HF|DD and pCAG-GFP-ATG13-1-230(HF|DD). All the plasmids were confirmed via sequencing and deposited to Addgene: pCAG-GFP-ATG13-1-230(HF|DD)(250344) pCAG-TSF-ULK1(249533); pCAG-TSF-ULK1-ADA (249534); pBMN-HA-ATG13-HF|DD (249535); pCAG-MBP-ULK1-1-829-mcherry(249729); pCAG-MBP-ULK1-1-290-mcherrry(249732); pCAG-MBP-ULK1-290-829-mcherry(249733); pCAG-ATG101ΔCTH(250340); pCAG-TSF-ATG101ΔCTH (WF|DD)(250341); pCAG-GST-ATG101(WF|DD)(250342); pCAG-GFP-ATG13-1-230(250343); pCAG-TSF-GFP-WIPI3 (257242); pCDH1-HA-ATG101-WT(257243); pCDH1-HA-ATG101-WF-DD (257244). All the constructs descripted above were listed in Table S1.

The following vectors were generated by MRC-PPU Reagents and Services, University of Dundee and are available on the website: pBABE puro mCherry-GFP-FIS 101-152 (*mito*-QC, DU55502), pBABE puro mCherry GFP LC3 (DU55696), pBabeD Hygro FLAG (DU37214), pBabeD hygro FLAG ULK1 (DU45617), pBabeD hygro FLAG ULK1 P433A V434D P435A (DU79458), pBabe puro-FLAG ATG13 siRNA resistant (DU79579) and pBabe puro-FLAG ATG13 H214D F215D siRNA resistant (DU79578). All the constructs descripted above were listed in Table S1.

### Protein expression and purification

Proteins that including GST-ATG13(6-196)/ATG101 and mutants, ATG13(1-230)-GFP/ATG101 and mutants, GST-WIPI1-4 and mutants, WIPI2d-TSF, TSF-FIP200(1-640)/ATG13/ATG101 and mutants, TSF-ULK1-WT and TSF-ULK1-WT-ADA were expressed in HEK293T-GNTI-cells. Variable volumes (0.3-1 L) cells were grown to a density of 2.0-2.5 x 10^6^ cells/L. 1 mg of total DNA/L were transfected using the polyethelenimine (PEI) (Polysciences) transfection method. For protein complexes, the plasmids were added in equal mass proportions such that the total DNA added equaled the final concentration of 1 mg DNA/L (e.g. transfection of 500 mL of cells with GFP-ATG13/TSF-ATG101 corresponded to 250 μg of pCAG ATG13 and 250 μg pCAG ATG101). After 48 hours, the cells were pelleted at 2200 RPM by centrifugation. The pellets were washed with PBS, spun using a tabletop centrifuge at 500 x g, flash frozen, and stored at -80 C until they were needed.

The Initial purification steps were roughly equivalent regardless of affinity tags. Cells were thawed to 4 C and resuspended in a buffer consisting of 25 mM (4-Hydroxyethyl)piperazine-1-ethanesulfonic acid (HEPES) pH 7.5, 200 mM NaCl, 2 mM MgCl2, 1 mM tris(2-carboxyethyl)phosphine (TCEP) and 10% Glycerol. An EDTA-Free Protease inhibitor tablet (Roche) was added, and the gently resuspended pellet was transferred to a Pyrex Dounce homogenizer. The cells were Dounce homogenized 30 times. The homogenate was transferred to a new tube and Triton X-100 was added to the cells for a final 1% concentration, gently mixed, and then left to rock at 4 C for 1 hour. Following detergent lysis, the cells were pelleted by centrifugation (17,000 x rpm for 45 minutes at 4 C).

Following pelleting, the supernatant containing the GST-tagged proteins were incubated with GSH resin pre-equilibrated with a buffer containing 25 mM HEPES, 200 mM NaCl, 2 mM MgCl_2_, 1 mM TCEP with gentle shaking for 2 hours. The protein-loaded resin was applied to a gravity column and washed extensively until the flow through was negative for protein, as assessed by Bradford reagent. The protein was eluted with a wash buffer containing 25 mM L-glutathione. The protein was loaded onto an S200 10/300 column (Cytiva) pre-equilibrated with 25 mM HEPES, 200 mM NaCl, 2 mM MgCl_2_, 1 mM TCEP. Peak fractions were pooled, flash frozen in liquid nitrogen and stored in 25 μL aliquots. For the WIPI3 used in GUV assays, following elution from the resin, the protein was cleaved overnight with TEV protease before application to an S75 10/300 column (Cytiva), and the peak fractions were pooled, concentrated, stored in 25 μL aliquots, flash frozen with liquid nitrogen, and stored at -80 C.

To purify tandem Strep-FLAG (TSF) tagged proteins, supernatant containing these proteins were applied to Strep-Tactin Sepharose (IBA biosciences) resin pre-equilibrated with a buffer containing 25 mM HEPES, 200 mM NaCl, 2 mM MgCl_2_, 1 mM TCEP at 4°C for 1-2 hours with gentle agitation. The resin loaded with protein was applied to a gravity flow column, and washed extensively until the flow through was free of protein, as assessed by Bradford reagent. For TSF-ULK1-WT and TSF-ULK1-ADA, the proteins were eluted with wash buffer containing 4mM desthiobiotin and 10% glycerol. The other TSF-tagged protein was eluted with wash buffer containing 1 mg/mL desthiobiotin. In the case of TSF-FIP200(1-640)/ATG13/ATG101, the protein was loaded onto an S6 10/300 column. Peak fractions were pooled, concentrated, flash frozen with liquid nitrogen, and stored at -80°C for downstream uses. GFP-ATG13(1-230)/TSF-ATG101, ATG13(1-230)/TSF-ATG101 were both applied to an S200 10/300 column (Cytiva) for size exclusion. Peak fractions were pooled, concentrated, flash-frozen with liquid nitrogen, and stored at -80°C for downstream uses.

To purify MBP tagged mCherry-ULK1^ΔEAT^ (1-828), mCherry-ULK1 ^ΔΚDΔEAT^ (291-828), and ULK1^KD^ (1-290) supernatant containing these proteins were applied to Amylose resin pre-equilibrated with a wash buffer containing 25 mM HEPES, 500 mM NaCl, 2 mM MgCl_2_, 1 mM TCEP at 4°C for 1-2 hours with gentle agitation. The resin loaded with protein was applied to a gravity flow column, and washed extensively until the flow through was free of protein, as assessed by Bradford reagent. The protein was eluted with wash buffer containing 40 mM amylose. The proteins were concentrated, frozen with liquid nitrogen, and stored at -80°C for downstream assays.

### AlphaFold Prediction pipeline and structure visualization

Multiple sequence alignments for ATG13(1-220) and ULK1 were generated using the DeepMSA2 server with the slow search settings (searched against Uniclust30, Uniref90, Metaclust, mGnify, BFD, TaraDB, MetaSourceDB and JGIclust). The resulting MSAs were used to generate the sequence logos in Figures 1 and Figures 3 using the weblogo tool(*65, 66*). The structural models of ATG13(1-220) + ATG101 + WIPI3 and ATG13(1-220) + ATG101 + ULK1(428-450) were predicted using the ColabFold server, with MSA generation using the default pipeline(*67–69*). The structures were analyzed and visualized in UCSF ChimeraX(*70*).

### GUV reconstitution and swelling

GUVs were prepared by hydrogel-assisted swelling similarly to as described in Chang et al, 2021(*71*). 100 μL of 5% (w/w) polyvinyl alcohol (PVA) with a molecular weight of 145,000 (Millipore) was coated onto a coverslip of 25 mm diameter that was cleaned sequentially with methanol, ethanol, and diH_2_O. The coated coverslip was placed in a heating incubator at 60°C to dry the PVA film for at least 30 minutes. For all GUV experiments, a lipid mixture with a molar composition of 64.9% 18:1 dioleoylphosphatidylcholine (DOPC), 20% 18:1 dioleoylphosphatidylethanolamine (DOPE), 10% 18:1 dioleoylphosphatidylserine (DOPS), 18:1 5% dioleoyloleoylphosphatidylinositol-3 phosphate (DOPI3P) and 0.1% Atto647-DOPE in chloroform at 1 mM concentration was spread uniformly onto the PVA film at a final lipid quantity of 50 nmol. The lipid-coated coverslip was then placed under vacuum overnight to evaporate residual chloroform. A 100 μL 400 mM sucrose solution (416 mOsm) was used for swelling at room temperature for 1 hour, and the GUVs were then harvested and used within 8 hours.

### GUV experiments

GUV experiments were conducted at room temperature in a LabTek 8 well glass observation chamber. First the chamber was passivated with 5 mg/mL β-casein for 0.5∼1 hour and then washed 3 times with a buffer containing 25 mM HEPES, 200 mM HEPES, 2 mM MgCl_2_, 1 mM TCEP. Reactions were prepared using 10 μL GUVs with or without 200 μM WIPI3, across concentrations of ATG13(1-230)-GFP/ATG101 including 50, 100, 200, and 500 nM, with three technical replicates. Images were acquired using a Nikon A1 confocal microscope under a 60 x oil objective. The fluorescence intensity was measured using a consistent laser power between technical replicates. Final signals were analyzed using Fiji. A line was drawn for every SUV with clear boundary and without aggregation, and the maximum intensity of GFP on the line was defined as the intensity of ATG13(1-230)/TSF-ATG101 enriched on GUV. Final statistics and curve graph were analyzed using Prism 10.

### SUV production

Liposomes were prepared by first drying overnight and rehydrating 0.43 mg of lipids (77% DOPC, 10%DOPE, 10% DOPS, and 3% PI3P) with 860 µL of buffer containing 25 mM HEPES (pH 7.5) and 200 mM NaCl on ice, resulting in a final concentration of 0.6 mM total lipid (0.5 mg/mL). The lipid suspension was vortexed for 5 minutes and subjected to 10 freeze-thaw cycles. The rehydrated lipids were then extruded 21 times through a 50 nm polycarbonate membrane using a mini extruder, producing 50 nm SUVs.

### Microscopy-based bead protein–protein interaction assay

Protein-protein interactions were assessed using Glutathione Sepharose bead-based pulldown assays. In Fig. 1F, GST itself, GST-WIPI3 (WT or mutants) and GST-WIPI4 were used at a concentration of 0.5 μM and immobilized on 5 μL of Glutathione Sepharose 4B beads (GE Healthcare) in reaction buffer containing 20 mM HEPES (pH 7.5), 200 mM NaCl, 2 mM MgCl_2_ and 1 mM TCEP for 5 min at room temperature. Then, GFP-ATG13(1-230)/ATG101 (WT or mutants) is added, immobilization was carried out at room temperature for 60 minutes, and the reaction mixes were applied immediately to a LabTek 8 well chamber for visualization. In Fig. 2, 1 µM GST-ATG13(6-196)/ATG101 (WT or mutant) was used as a bait to incubate with 5 µL GSH beads and 100 µM Atto647 LUV. In Fig. 5, GST-GFP-ULK1(415-464) (WT or ADA mutant) acted as a bait to incubate with GSH beads and mcherry-ATG13(1-230)/TSF-ATG101. And for Fig, S3, GST itself or GST-ATG13(6-196)/ATG101 acted as a bait to incubate with MBP-mCherry-ULK1(1-828), MBP-mCherry-ULK1(291-828) or MBP-mCherry-ULK1(1-290) separately. Images were acquired using a Nikon A1 microscope. Views were selected randomly, and images were acquired for single time points across the replicates.

### Membrane reconstitution assay

Reactions were carried out in a total volume of 18 µL, composed of 6 µL of each sample from three experimental groups, initially without ATP. Reactions contained 3 nM ULK1+/-, 30 nM FIP200/ATG13 (WT or HF|DD)/ATG101 (WT or WF|DD), 100 nM WIPI3 +/-, 100 nM WIPI2 +/-, 100 nM ATG16(78-300), or 200 μM LUVs +/-. Samples were incubated at room temperature (RT) for 10 minutes, followed by cooling on ice. Reactions were initiated by the addition of ATP and incubation at 30 °C. Aliquots of 3 µL were taken at 10-, 30-, and 60-minutes post-initiation, and each was immediately quenched by the addition of 12 µL of 8 M urea. Aliquots (2 µL) of each denatured and diluted sample were applied onto a nitrocellulose membrane and allowed to dry for 10 minutes. Membranes were blocked overnight at 4 °C with 3% milk in TBST. After washing, membranes were incubated with primary rabbit anti-ATG16L1 phospho-S278 antibody (Rb pATG16L1-S278, 1:5000 dilution) at RT for 1 hour. Following additional washes, membranes were incubated with HRP-conjugated anti-rabbit secondary antibody (1:5000 dilution) for 50 minutes at RT. Detection was performed using the Thermo Scientific SuperSignal™ West Femto Maximum Sensitivity Substrate (0.5 mL + 0.5 mL mixed components).

### Halo assay to assess autophagy

In the first day, Halo-LC3, ATG13-KO+Halo-LC3, ATG13-KO+Halo-LC3+HA-ATG13-WT and ATG13-KO+Halo-LC3+HA-ATG13-(HF|DD) stable cell lines were seeded into 24 well plate at 5x10^4 per well. Next day, change to the fresh medium 2 hours before adding 100nM Janelia Fluor® 549 HaloTag® Ligand. After treatment with Halo ligand for 30min, washed with PBS for 2 times. Then, treated with EBSS for indicated times. Finally, lysis the cell with 2% SDS extraction buffer ((50mM Tris pH 6.8, 2% SDS, 1% mercaptoethanol, 12.5% glycerol, 0.04% bromophenol blue, and the protease inhibitor cocktail (Roche, 04693132001)). Proteins were separated using a 4-12% Bis-Tris SDS-PAGE (Invitrogen) and detected the fluorescent signal of Halo-ligand in gel.

### Western blot and co-immunoprecipitation (Co-IP)

Cells were treated and proteins were extracted as described above. After separation with 4-12% Bis-Tris SDS-PAGE or 6–14% Bis Tris gels, immunoblotting was performed with indicated primary and second antibodies.

To detect the interaction of ATG13 with WIPI3 in cells, ATG13-KO+Halo-LC3, ATG13-KO+Halo-LC3+HA-ATG13-WT, and ATG13-KO+Halo-LC3+HA-ATG13-(HF|DD) stable cell lines were seeded into 10 cm dishes at 1×10⁵ cells/ml. The next day, 10 μg of pCAG-TSF-GFP-WIPI3 plasmid was transfected into each group using Lipofectamine 2000 (Thermo Fisher, cat. #11668019) at a DNA:lipid ratio of 1:2 (μg:μl) for 24 h. Medium was then replaced with fresh medium 2 h before treatment with EBSS for 1 h. Cells were harvested with pre-chilled Co-IP lysis buffer (50 mM Tris-HCl, pH 7.5, 200 mM NaCl, 10 mM EDTA, 5% glycerol, 1% NP-40, supplemented with protease inhibitor cocktail). After lysis for 1 h at 4°C and centrifugation at 13,000 × g for 10 min, 1/20 of the supernatant was saved as input, and the remainder was incubated with Pierce™ Anti-HA Magnetic Beads (Thermo, 88836) at 4°C overnight. The bead–protein complexes were then washed five times with Co-IP lysis buffer, 10 min per wash. Finally, bound proteins were eluted by boiling in 2× SDS loading buffer for 10 min before being resolved by SDS-PAGE.

To detect the interaction between FLAG ULK1 and endogenousATG13, cell lysates were collected using NP-40 lysis buffer supplemented with protease inhibitor cocktail and phosphatase inhibitor cocktail and protein concentration was determined using BCA (Pierce BCA Protein Assays). For FLAG pulldown, 0.6 mg of lysate was incubated with 30 μl anti-FLAG M2 affinity beads (MRC-PPU Reagents and Services) for 4 h at 4°C in a rotor (25 rpm). Following incubation beads were precipitated by centrifugation and washed 3 times with NP-40 lysis buffer. Proteins were eluted using 40 μl of 2X LDS (Life technologies, #NP0007) +1mM DTT and 1 μl of Beta-mercaptoethanol, and heat at 95°C for 5 min. Eluates were passed through a Spin-X® (Sigma, #CLS8162) column to separate the beads.

### Antibodies and reagents

Primary antibodies used were as follows: anti-HIF1α (R&D system MAB1536, 1:1,000 for WB), anti-HSP60 (CST #4870S, 1:1,000 for WB), anti-OMI (MRC PPU Regeants and Services, University of Dundee, 1:500 for WB), anti-vinculin (Abcam ab129002, 1:10,000 for WB), anti-COXIV (CST #4850S, 1:1,000 for WB), anti-FLAG HRP (Sigma, A8592, 1:2,000 for WB), anti-FLAG (Sigma, F1804, 1:500 for IF and WB), anti-Phospho-Atg14 (Ser29) (CST #92340S, 1:1,000 for WB), anti-Atg14 (CST #96752S, 1:1,000 for WB), anti-FIP200 (Proteintech, 17250-1-AP, 1:500 for IF and 1:1,000 for WB). Rabbit p-ATG16L1(S278) (Abcam, ab195242, 1:1,000 for dot blot and CST #45511S, 1:1000 for WB), Rabbit anti-ATG16L1 (Abcam ab187671, 1:1000 for WB), Rabbit anti-ATG13 (CST, 13468, 1:1,000 for WB and Sigma SAB4200100, 1:1,000 for WB), anti-ULK1 (CST #8054S, 1:1,000 for WB), anti-ATG101 (Sigma SAB4200175, 1:500 for WB), Mouse anti-GAPDH (Abcam, ab8245, 1:5000 for WB). Rabbit anti-HA (CST, 3724, 1:200 for IF), Secondary antibodies used were as follows: goat anti-Rabbit IgG (H+L) HRP conjugate, goat anti-mouse IgG (H+L) HRP conjugate and Donkey anti-sheep IgG (H+L) HRP conjugate were purchased from Thermo Scientific (#31460, #31430, #A16041, respectively, 1:5,000 for WB).goat antirabbit-Alexa Fluor 568 (Life Technologies, A11012, 1:1000 for IF).

3-Hydroxy-1,2-dimethyl-4(1H)-pyridone (DFP) (Sigma 379409), Ponceau S (Sigma P3504). Earle’s Balanced Salt Solution (EBSS, Gibco 14155). All the agents descripted above were listed in Table S1.

### Immunofluorescence

For the results in Fig.6, cells were seeded onto sterile glass coverslips in 24-well dishes. Coverslips were washed once with PBS, fixed with 3.7% (w/v) formaldehyde, 200 mM HEPES pH 7.0 for 10 min and washed twice with PBS. Cells were permeabilized with 0.1% triton in PBS for 5 min. After two washes with PBS, samples were blocked by incubation for 30 min in blocking buffer (1% (w/v) BSA/PBS). Coverslips were incubated for 2 h at 37°C with primary antibodies in blocking buffer and washed three times in PBS. Coverslips were then incubated for 1 h at room temperature with Alexafluor coupled secondary antibodies (Life Technologies) in blocking buffer and washed an additional three times in PBS. If needed, nuclei were counterstained with 1 μg/ml Hoechst-33258 dye (Sigma) for 5 min, and slides were washed twice with PBS and mounted in ProLong Gold (Invitrogen). Observations were made with Nikon Eclipse Ti2-E microscope (CFI Plan Apochromat Lambda D 60X Oil objective). Images were processed using Fiji (ImageJ) and Adobe Photoshop Software.

### Modeling and simulation of ATG13-ATG101-WIPI3 and ATG13-ATG101-WIPI3-WIPI2 complex

The structure of the ATG13-ATG101-WIPI3 and ATG13-ATG101-WIPI3-WIPI2 complexes were modeled with AlphaFold v2.3.0(*67*). To associate the protein complex with the membrane, we initially docked a PI3P lipid near the LRRG motif of WIPI3 (FRRG motif of WIPI2 in the case of ATG13-ATG101-WIPI3-WIPI2 complex), positioning the P3 atom of PI3P at a distance of ∼2.9 Å from R225. Next, we prepared an atomistic membrane bilayer of area 13.3 nm ×13.3 nm consisting of 65 % DOPC, 20 % DOPE, 10 % DOPS, and 5 % PI3P using CHARMM-GUI(*72, 73*). We then placed the PI3P-bound ATG13-ATG101-WIPI3 complex and, in a separate setup, the ATG13-ATG101-WIPI3-WIPI2 complex onto the membrane, ensuring that the WF finger and ATG101’s C-terminal amphipathic helix (G201-L218) remained inserted in the bilayer. The initially docked PI3P lipid also remained embedded in the bilayer. To remove steric clashes and yield efficient protein insertion into the membrane, we removed a total of 47 (51 in the case of ATG13-ATG101-WIPI3-WIPI2) lipids near the WF finger, the C-terminal amphipathic helix and the initially docked PI3P. We then solvated the system with TIP3P water and maintained a physiological salt concentration (150 mM NaCl). Subsequently, we carried out energy minimization, equilibration, and production runs with this solvated and ionized structure employing GROMACS 2021.5(*74*) and the CHARMM36m(*75*) force field. Minimization for 5000 steepest descent steps was followed by MD equilibration first under NVT (250 ps) and then under NPT (∼2 ns) condition. We applied harmonic position restraints on the heavy atoms of the protein during the equilibration simulations which were gradually lifted. In the first 125 ps, the protein backbone and sidechains were restrained with force constants of 4000 and 2000 kJ·mol-1·nm-2, respectively, using a Berendsen thermostat(*76*) and a 1 fs integration timestep. To maintain membrane bilayer integrity, harmonic position restraints were applied exclusively along the Z-axis (perpendicular to the membrane plane) of the headgroup of the lipid molecules using a force constant of 1000 kJ·mol-1·nm-2. In the next 125 ps of equilibration, the force constants on the protein backbone, sidechain, and the lipid head groups were reduced to 2000, 1000, and 400 kJ·mol-1·nm-2, respectively. Subsequently, a semi-isotropic pressure coupling was activated using a Berendsen barostat and the force constants of the harmonic restraints were further reduced to 1000 and 500 kJ·mol-1·nm-2 on the protein backbone and sidechain, respectively, over 125 ps while keeping the restraints on the lipid headgroups intact. Next, the time step of integration was increased to 2 fs and the force constants of harmonic position restraints were further reduced to 200, 50, and 40 kJ·mol-1·nm-2 on the protein backbone, sidechain, and lipid headgroups, respectively, during a 1 ns equilibration phase. Finally, the system was equilibrated for an additional 1 ns with only minimal protein backbone restraints (50 kJ·mol-1·nm-2) to allow full relaxation before unrestrained production simulation. The equilibrated system was subjected to production runs in the NPT ensemble for 1 μs. To obtain statistically reliable insights into the dynamics, four independent replica simulations were carried out. No position restraints were appplied during the production runs. The integration timestep was 2 fs. For the ATG13–ATG101– WIPI3–WIPI2 complex, an additional ∼250 ns equilibration was performed prior to the production run. During this phase, 18 harmonic distance restraints were applied to key molecular interfaces (ATG13:WIPI3, ATG13:WIPI2, the LRRG/FRRG motifs of WIPI3/WIPI2, and PI3P) to ensure full relaxation of the complex, its IDRs, and the interacting lipids and solvent in the subsequent unrestrained simulation. The system temperature and pressure were maintained at 303 K and 1 bar, respectively, using the Nosé-Hoover thermostat(*77, 78*) and a semi-isotropic Parrinello-Rahman barostat(*79*). Long range electrostatic interactions were treated using the particle-mesh Ewald (PME)(*80*) algorithm. The LINCS algorithm(*81*) was used to constrain covalent bonds involving hydrogen atoms. Simulation trajectories were analyzed with VMD(*82*). For the salt-bridge distance analysis, we used the OD1 and OD2 atoms of Asp and the NZ atom of Lys. The RMSD evaluation focused on backbone atoms of the DHF motif and proximal WIPI3 residues (within 5 Å: V35, K44, R62 C63 N64 D87 and L88).

### Modeling of ULK1C

We used a combination of computational approaches to generate a structural model of ULK1C tethered to the phagophore membrane. To do this, we combined a publicly available implementation of hierarchical chain growth (HCG)(*83, 84*) (https://bio-phys.pages.mpcdf.de/hcg-from-library/) with a cyclic coordinate descent (CCD) algorithm(*85*). HCG is a fragment-assembly mechanism that produces structural ensembles of IDRs in good agreement with experimental probes of both local and global structure. In CCD, we model a disordered region of a protein as a kinematic chain with rotatable bonds acting as hinges. In this implementation, we iteratively draw a random residue on the chain modelled by HCG and a ϕ or ψ dihedral angle, then perform a small rotation drawn from a Gaussian distribution with a mean of zero and a standard deviation of 1 degree. We accept the move if the distance between the end of the kinematic chain, the effector arm, and some target is reduced, and no clashes are created. The effector arm can be a residue on either terminus or a point on an attached rigid body.

Starting from a simulation snapshot of the ATG13-ATG101-WIPI3-WIPI2 complex on the membrane, we first generated a structural ensemble of the disordered region of ATG13 (residues 230 to 363) using HCG. For any HCG sampling, we only accept conformations that place the IDR itself and any attachments fully above the membrane and do not create clashes when added to the complex. The ULK1C core was modeled using AlphaFold2 and attached to HCG models as a rigid body via alignment on overlapping residues. We then selected IDR conformations that placed C927 and C1003 close to the membrane and performed CCD on the disordered region to minimize the RMSD of cysteine sulfur atoms to the membrane. We included a third point on the FIP200 arm since it has to lie flat on the bilayer and we do not check for clashes with the membrane during CCD minimization. The process was terminated when all three atoms were within 5 Å of the approximate height of the lipid headgroups. From these doubly membrane-bound conformations, we then sampled the ULK1 IDR (residues 277 to 831) in the absence of binding to the ATG13 HORMA domain using HCG with an AlphaFold2 model of the ULK1 KD attached. To account for the binding, we modeled ULK1 residues 428 to 450 bound to ATG13 using AlphaFold2 and placed it into our complex structures via superposition of the ATG13 HORMA domain. Then, we selected ULK1-IDR conformations close to ATG13, removed residues 1 to 449 and minimized the alignment RMSD of the overlapping residue using CCD. The iteration was terminated when the alignment RMSD along backbone heavy atoms was below 0.6 Å. Lastly, we resampled the remaining N-terminal residues (277 to 428) with the KD attached. To gauge the effect of ULK1-IDR binding to the ATG13 HORMA domain, we measured the distance of the KD center of mass to the membrane with and without including the bound segment. We report distances for three separate iterations of the procedure starting from different initial conformations of ATG13.

### Quantification of Immunofluorescence image

#### 1 Quantification for beads binding assay

To quantify protein binding, we developed a semi-automated pipeline, beads were first identified in the brightfield channel using Canny edge detection followed by a Hough Circle Transform. To ensure data quality, an overlap filter was applied to exclude clustered beads. The detected coordinates were then mapped onto the GFP or RFP channel, where a disk mask (optimized via a radius-refinement step) was used to quantify the mean fluorescence intensity of the protein bound to each bead. The automated pipeline ensures unbiased quantification even for weakly binding mutants, as the bead detection is performed independently in the brightfield channel, avoiding any ’picking bias’ in low-signal GFP images. The code for quantification has been deposited in Github: https://doi.org/10.5281/zenodo.20076398.

#### 2 ULK1 puncta

A semi-automated method was used to quantify ULK1 puncta. For each biological replicate, at least 70 cells were analyzed per condition. Images were processed in **Fiji,** where a threshold mask was applied to each fluorescence channel using identical threshold values across all images within a replicate. Individual cells were manually outlined and added to the regions of interest (ROIs). The number of puncta above the threshold was then automatically quantified using the *Analyze Particles* function.

### Autophagy and Mitophagy assay

Cells stably expressing mCherry-GFP-LC3B or mito-*QC* mitophagy reporter system (mCherry-GFP-FIS1 101–152) were seeded onto sterile glass coverslips in 24-well dishes. After treatment, coverslips were washed twice with PBS, fixed with 3.7% (w/v) formaldehyde, 200 mM HEPES pH 7.0 for 10 min and washed twice with PBS. After nuclei counterstaining with 1 μg/ml Hoechst-33258 dye, slides were washed and mounted in ProLong Gold (Invitrogen).

### Flow cytometry analysis

A total of 1.75 × 10⁵ cells were seeded in a 6 cm dish. Following treatment, cells were washed with PBS, trypsinized for 5 min, neutralized with PBS containing 2% FBS, and centrifuged at 1,200 rpm for 3 min. The pellet of cells was resuspended in 250 μl of PBS and 1 ml of 3.7% (w/v) formaldehyde, 200 mM HEPES pH 7.0 were added. After 30 min at RT, 3 ml of PBS+2%FBS was added before centrifugation 5 min at 1,200 rpm. Finally, the pellet of cells was resuspended in 2% FCS in PBS and analyzed by flow cytometry. For each independent experiment, at least 50 000 events were acquired on LSRFortessa cell analyzer. Based on FCS and SSC profiles, living cells were gated. As negative control, cells expressing any autophagy or mitophagy reporter were used. To quantify the percentage of cells undergoing autophagy or mitophagy, the ratio GFP/mCherry was analyzed. The gate used for the nontreated condition or control cells was applied to all the other conditions.

### Statistical analysis

Representative results of at least three independent experiments (biological replicates) are shown. GraphPad Prism software was used for all statistical analyses. In Fig 1G, One-way ANOVA+ Bonferroni post-hoc test was used. In Fig.2C, 4C, 4E, 4G, 4J, 4L,5G, 6B, 6D, 6F, 6H, S2C, S6B, S10B and S10D, the statistical significance was determined by two-way ANOVA with Sidak’s multiple comparisons test or Tukey’s multiple comparisons test. Otherwise, the unpaired, two-tailed Student’s t-test was used. P-values are indicated as *P < 0.05, **P < 0.01, ***P < 0.001 and ****P < 0.0001. ns: P > 0.05. The error bars represent the standard deviations (s.d.) or standard error of the mean (SEM) as indicated in figure legends.

## Acknowledgements

We thank D. Fracchiolla and Pei-I Tsai for program management support, and M. Lazarou and all members of ASAP team mito911 for advice and discussions.

## Author Contributions

Y.D., S.P., L.P.W., I.G., and J.H.H., Writing-original draft; S.P., X.R., I.G., G.H. and J.H.H., Conceptualization; Y.D., Y.L, S.P., J.B., L.P.W., A.S.I.C., X.R., and E.A., Investigation; Y.D., S.P., J.B., L.P.W., A.S.I.C., E.A., I.G., G.H. and J.H.H., writing-review & editing; Y.D., S.P., J.B., L.P.W., X.R., and I.G., Methodology; Y.D., S.P., X.R., S.M., I.G., and G.H., Resources; Y.D., S.P., and A.S.I.C., Data curation; Y.D., S.P., J.B., L.P.W., A.S.I.C., and X.R., Validation; Y.D., Y.L, S.P., J.B., L.P.W., and A.S.I.C., Formal analysis; S.P., J.B., and L.P.W., Software; Y.D., S.P., J.B., L.P.W., and A.S.I.C., Visualization; S.M., I.G., G.H. and J.H.H. Funding acquisition; I.G., G.H. and J.H.H. Supervision; I.G., G.H. and J.H.H., Project administration.

## Data, Code, and Materials Availability

All data are provided in the manuscript or the source data associated with the manuscript (Table S1). The plasmids are available by request at Addgene.org with no restrictions beyond those of the UBMTA. Requests for the cell lines should be submitted to: IGG, or JHH. All data cleaning, preprocessing, analysis, and visualization was performed using Graphpad Prism (Ver.10).

**Figure S1.**
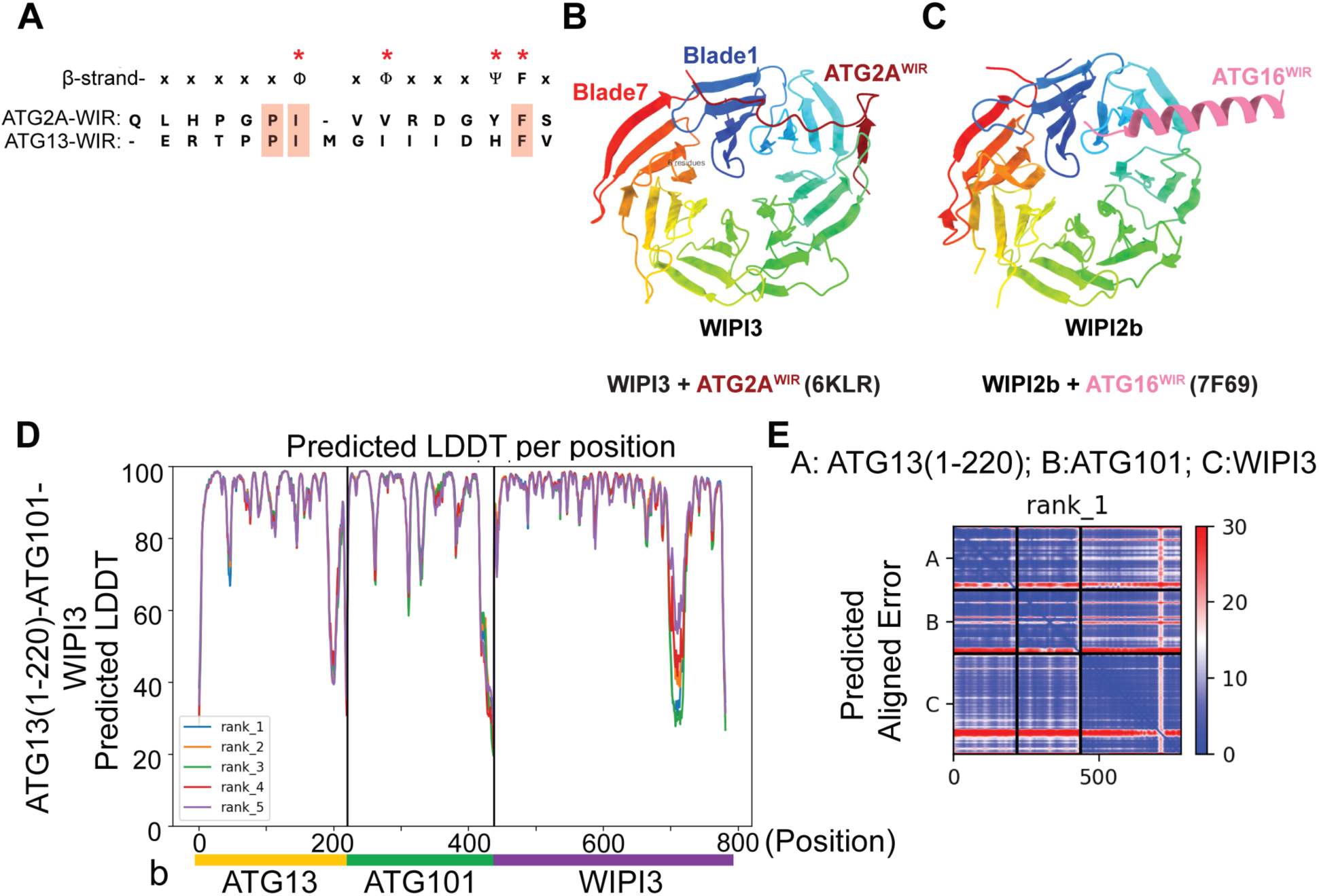
Comparison of the predicted binding between WIPI effectors and Structure prediction confidence of ATG13-ATG101-WIPI3. **(A)** Comparison of human ATG2-WIR and ATG13-WIR. Φ is a hydrophobic residue and Ψ is an aromatic residue. **(B)** Crystal structure (PDB: 6KLR) of the ATG2A WIR motif bound to WIPI3 in the pockets between blades1 and 2, and blades2 and 3. **(C)** Crystal structure (PDB: 7F69) of ATG16 W2IR motif bound to WIPI2 at the cognate blade2 and 3 pocket of WIPI2. (**D, E**) pLDDT (D) and PAE (E) of Alphafold model of ATG13-ATG101-WIPI3 in Fig. 1C. Colab AlphaFold provided 5 models (rank_1-rank_5), rank_1 model was chosen to show in Fig. 1C.

**Figure S2:**
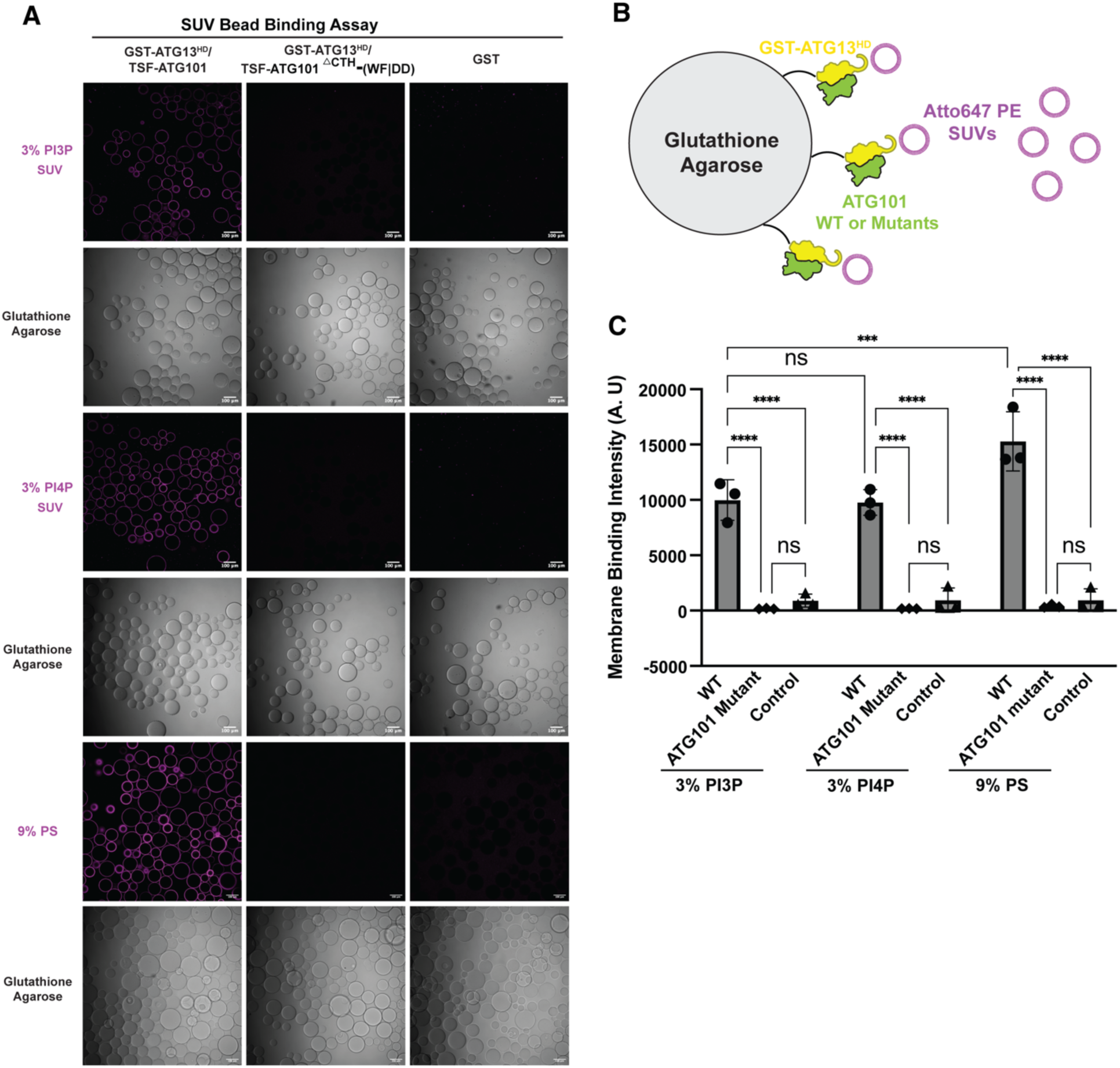
ATG13:ATG101 HORMA recruited to high curvature membrane by SUV-bead binding assay. **(A)** Fluorescent images show the results of GST beads binding experiments between GST-ATG13/ATG101 HORMA and SUVs with 3% PI3P, 3% PI4P or 9% PS. The scale bar is 100 µm. **(B)** Schematic of SUV and GST bead binding assay. ATG13:ATG101 HD was included at 1 μM concentration, and lipids were included at 100 µM. **(C)** Quantification of SUV binding to glutathione agarose beads as recruited by GST tagged ATG13:ATG101 WT or ATG13:ATG101 mutant. Three biological replicates were performed, with at least three images taken for each replicate. The average fluorescence intensity of all beads in each image was calculated. ***P<0.001, ****P<0.0001, ns, non-significance.

**Figure S3.**
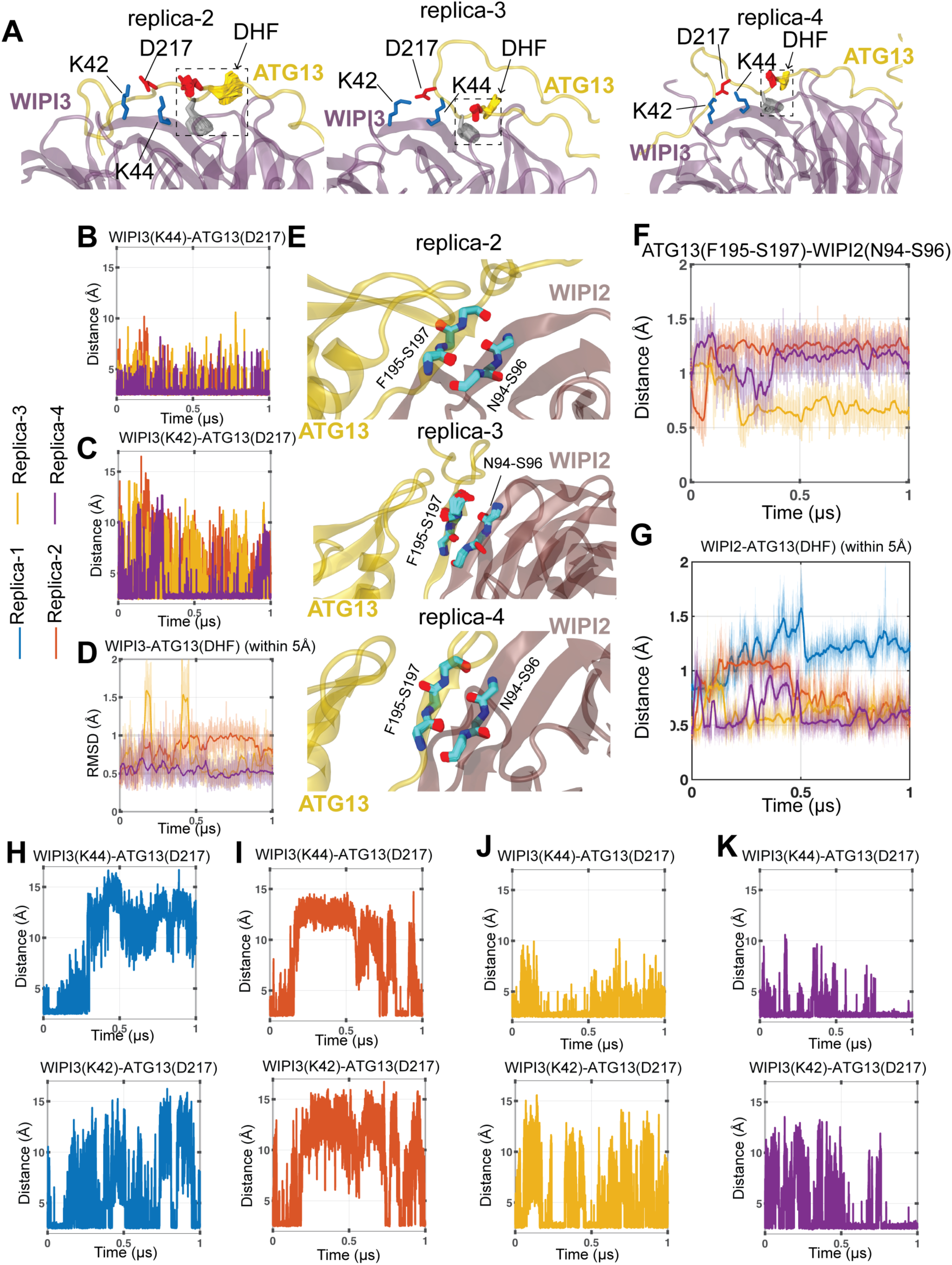
Dynamics of membrane-bound ATG13–ATG101–WIPI3 and ATG13–ATG101– WIPI3–WIPI2 complexes across multiple replica simulations. **(A)** Zoomed-in view of the ATG13–WIPI3 interface (as shown in Fig. 3B for replica 1) in replicas 2, 3, and 4 of the ATG13–ATG101–WIPI3 simulation. **(B, C)** Time evolution of the minimum distance between the OD1/OD2 atoms of ATG13 D217 and the NZ atom of WIPI3 K44 (B) or K42 (C) in replicas 2–4 (replica 1 shown in Fig. 3C, 3D). **(D)** Backbone RMSD of the ATG13 DHF motif and WIPI3 residues within 5 Å of this region for replicas 2–4 (replica 1 shown in Fig. 3E). **(E)** Zoomed-in views of the ATG13–WIPI2 interface in replicas 2, 3, and 4 of the ATG13– ATG101–WIPI3–WIPI2 simulation. **(F)** Backbone RMSD of the residues shown in (F) as a function of simulation time across different replicas (replica 1 shown in Fig. 3I). **(G)** Backbone RMSD evolution of the ATG13–WIPI3 interface (as defined in D for ATG13-ATG101-WIPI3) in different replicas of the ATG13–ATG101–WIPI3–WIPI2 simulation. **(H-K)** Time evolution of the minimum distance between the OD1/OD2 atoms of ATG13 D217 and the NZ atoms of WIPI3 K44 and K42 in replicas 1 (H), 2 (I), 3 (J), and 4 (K) of the ATG13–ATG101–WIPI3–WIPI2 simulation.

**Figure S4:**
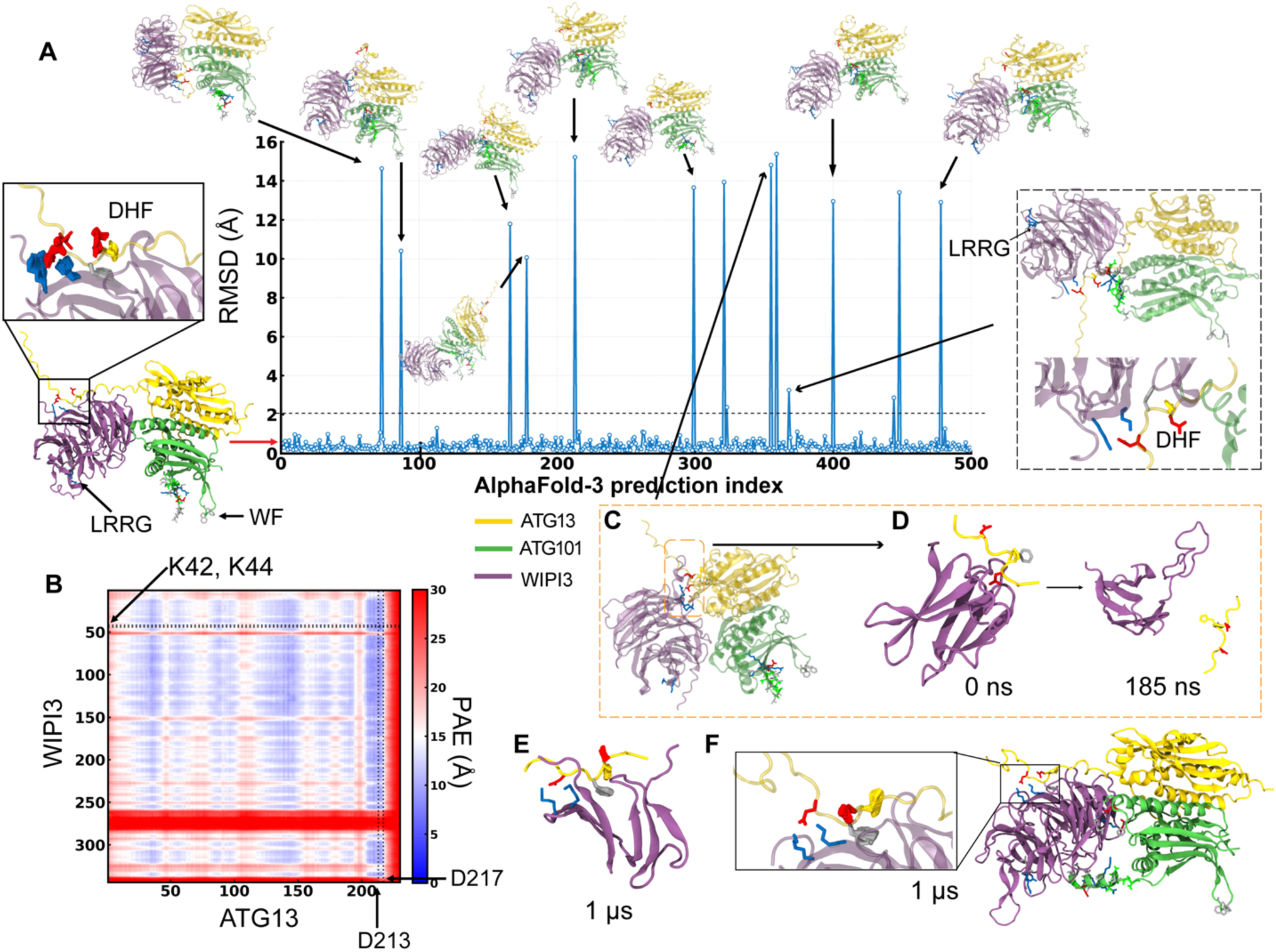
Structural validation of the ATG13-ATG101-WIPI3 interface using multiple AlphaFold-3 iterations and stability analysis of ATG13-WIPI3 interface fragments using MD simulations. **(A)** RMSD analysis of the ATG13-WIPI3 interface relative to the reference structure in Fig. 3. Key interface residues—including the ATG13 DHF motif and D217, the WIPI3 K42–K44 residues and LRRG motif, and the ATG101 WF finger—are rendered as sticks. The inset displays the superimposed coordinates of the DHF motif for all models within this primary cluster. Outlier structures (RMSD > 2 Å) are presented. The model within the black box exhibits a conserved ATG13-WIPI3 interface (zoomed in view below) but features a flipped LRRG motif relative to the WF finger. **(B)** PAE scores for the WIPI3-ATG13 interface, demonstrating high-confidence predictions centered on the DHF motif. **(C, D)** A specific outlier conformation model in A (C) and its interface fragment (D). Dissociation of this fragment occurred within 185 ns. **(E)** The ATG13-WIPI3 interface from the dominant cluster remains stable throughout the 1 *μs* simulation; sidechains of the DHF motif are shown superimposed across all trajectory frames. **(F)** Stability of the full hetero-trimeric complex from the dominant cluster, highlighting the preserved D217– K42/K44 salt bridge and stable binding of the DHF motif during the 1 *μs* membrane-free simulation.

**Figure S5:**
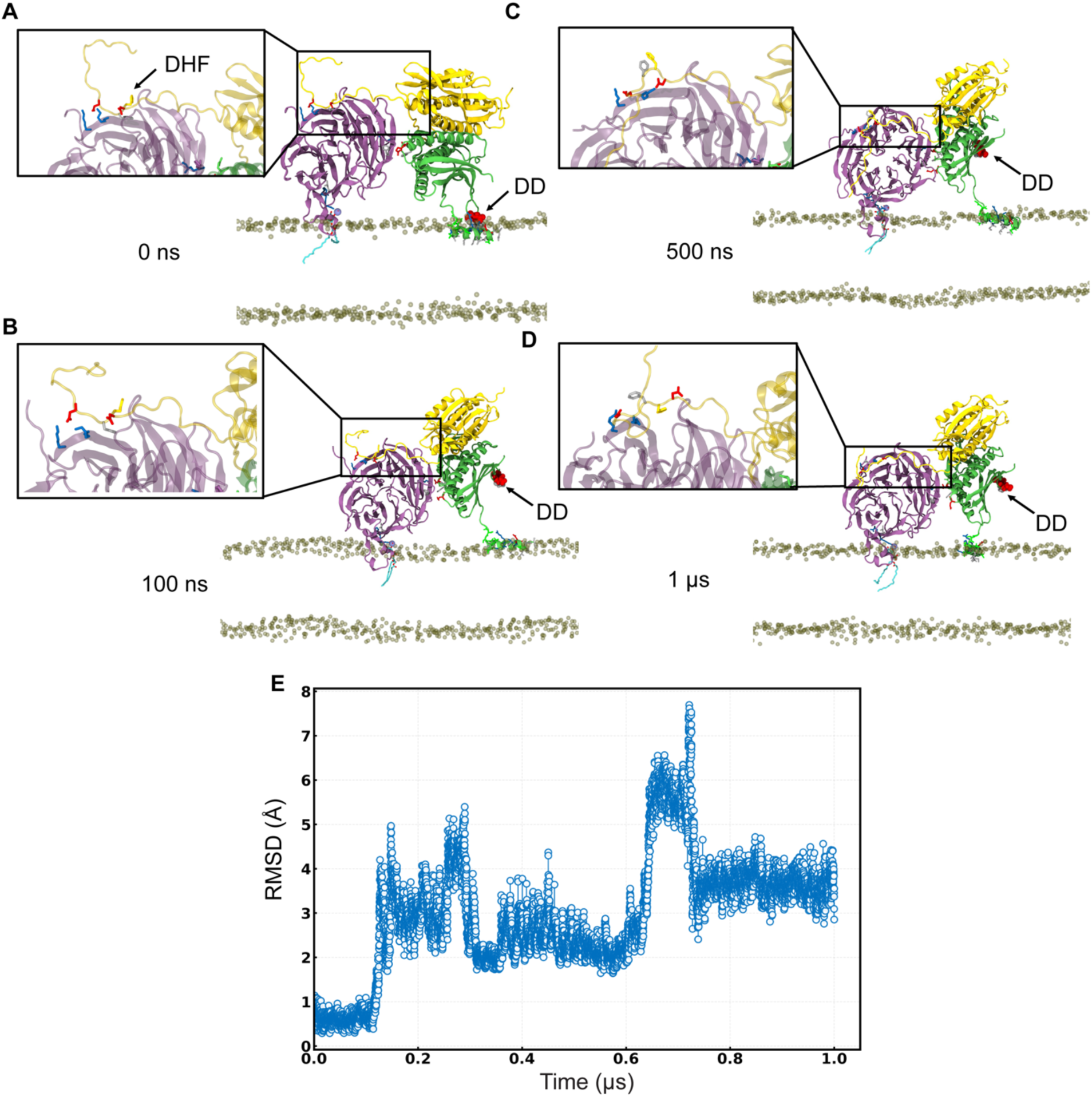
Impact of the ATG101 WF-to-DD mutation on the stability and membrane association of the ATG13-ATG101-WIPI3 complex. **(A-D)** Snapshot of the complex in the presence of the membrane (only phosphate groups are shown for clarity) at (A) 0 ns (B) 100 ns (C) 500 ns and (D) 1 *μs*. The newly introduced DD finger of ATG101 reverts to a solvent-exposed state away from the membrane. Insets show a magnified view of the ATG13-WIPI3 interface, with key residues (DHF motif and D217 of ATG13; K42/K44 of WIPI3) shown as sticks. **(E)** RMSD evolution of the DHF motif of ATG13 and its nearby WIPI3 residues exhibit a significant destabilization of the interface due to the WF to DD mutation.

**Figure S6:**
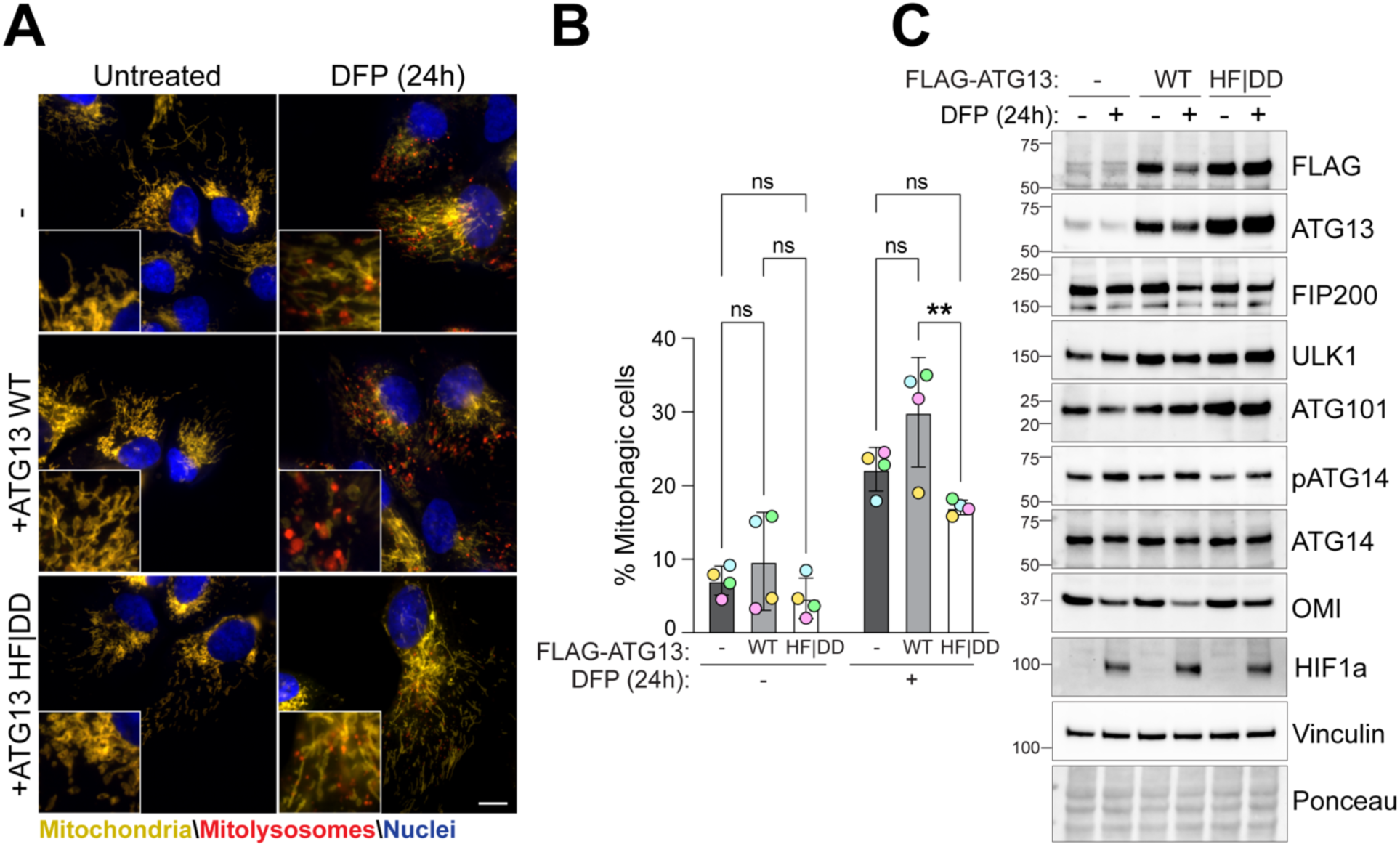
Effect of ATG13 HF|DD mutation on mitophagy. (**A**) Representative wide-field images of ARPE-19 cells stably expressing WT or HF|DD mutant FLAG tagged ATG13, and the *mito*-QC reporter. Cells were treated with 1mM DFP for 24 hours prior to imaging. (**B**) Flow cytometry analysis of the mCherry/GFP ratio (n = 4 biological replicates); statistics: two-way ANOVA with Tukey’s multiple-comparisons test. **(C)** Representative immunoblots of the indicated proteins from cells as in (A). **P<0.01, ns, non-significance.

**Figure S7:**
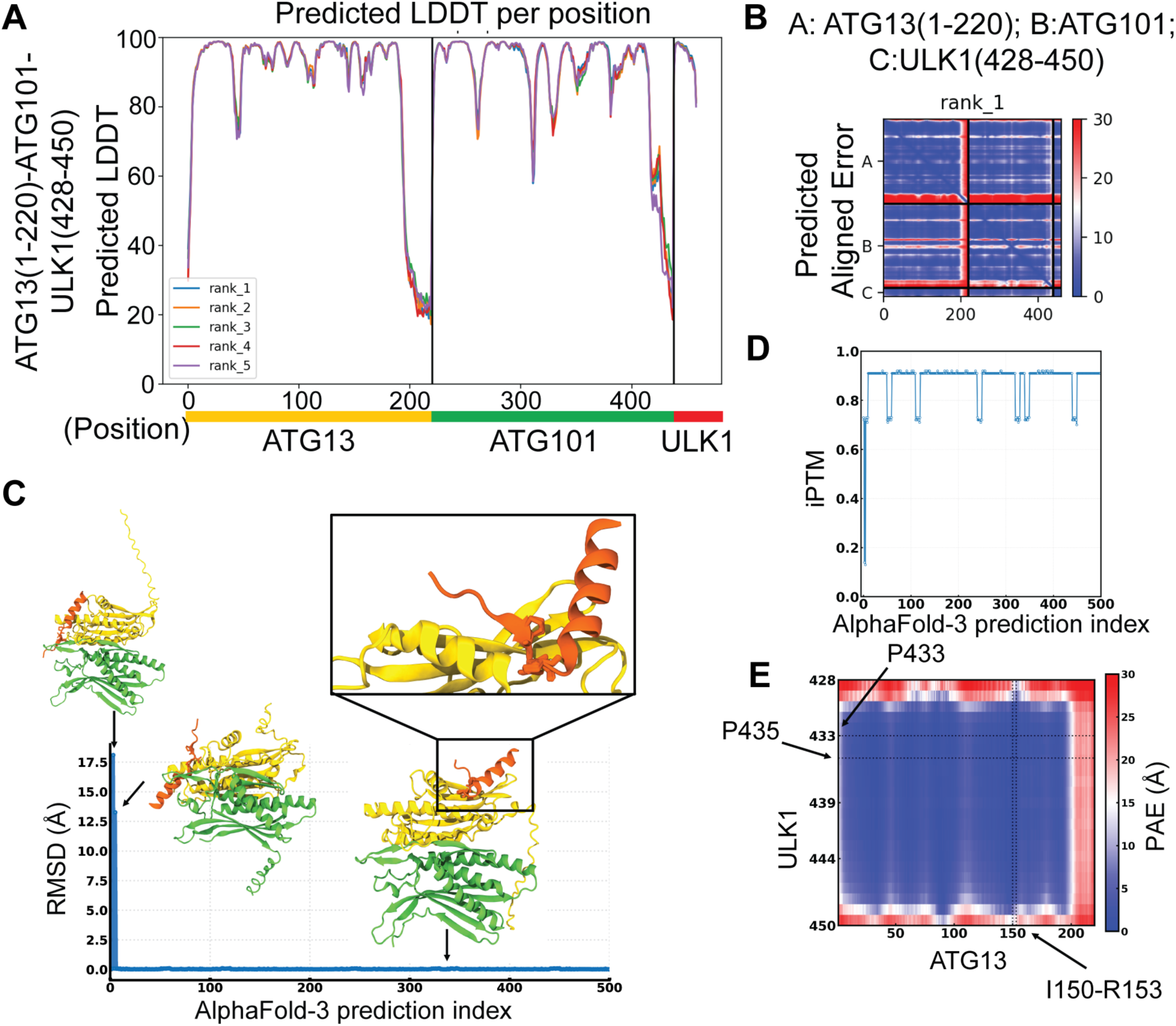
Structure prediction confidence of ATG13-ATG101-ULK1. **(A, B)** pLDDT (A) and PAE (B) of Alphafold model of ATG13-ATG101-ULK1 in Fig. 5C. Colab AlphaFold provided 5 models (rank_1-rank_5), rank_1 model was chosen to show in Fig. 5C. **(C)** RMSD analysis of the ULK1 (residues 428–450)–ATG13 (residues 150–153) interface relative to the reference structure in Fig. 5C identifies only two outliers (representative snapshot shown) among 500 total predictions. A snapshot of the dominant prediction is shown, with key ULK1 residues (P433, V434, P435) displayed as sticks. The inset shows the superposition of the PVP motif coordinates across all models within the primary cluster. **(D)** iPTM scores for all predicted ULK1 IDR–ATG13 models show consistently high values, with the exception of the outliers. **(E)** PAE score of a representative ULK1 IDR–ATG13 interface indicates high confidence, reflected by low PAE values. ULK1 residues P433 and P435 are highlighted with dotted lines, showing contacts with ATG13 residues I150–R153 (indicated by a dotted vertical line).

**Figure S8.**
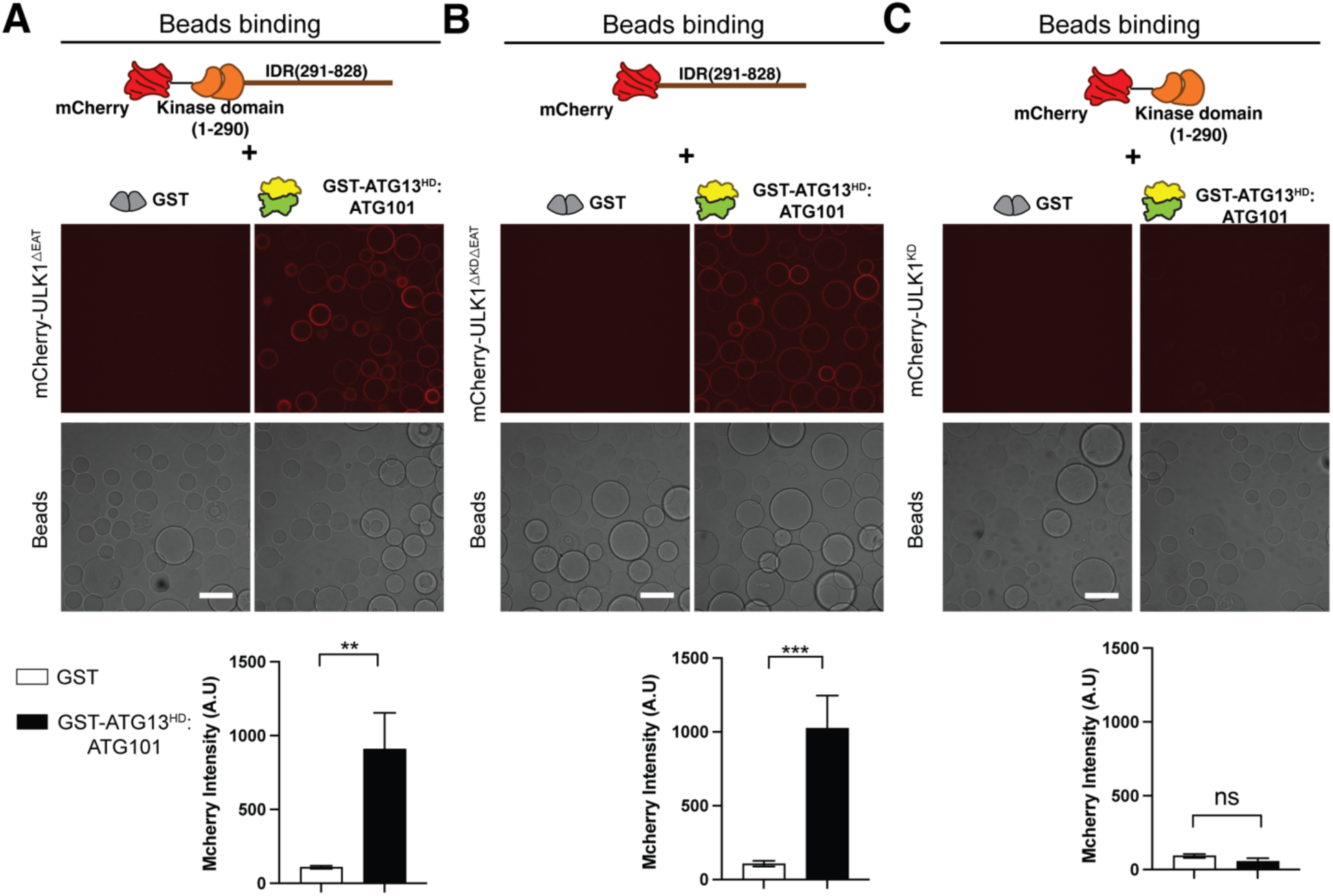
The IDR of ULK1 binds to ATG13(HORMA)/ATG101 directly. **(A)** ULK1 constructs lacking the EAT domain can bind to GST-ATG13^HD^:ATG101, the bar graph in below showed the quantification of intensity of mcherry channel per bead; **(B)** ULK1-IDR maintain the ability that bind to GST-ATG13^HD^:ATG101, the quantification showed in below. **(C)** ULK1^KD^ lost its binding to GST-ATG13^HD^:ATG101. n=3 biological replicates, statistics: Unpaired t-test. Scale bar is 100 µm. Mean ± SEM was shown. **P<0.01, ***P<0.001, ns, non-significance.

**Figure S9:**
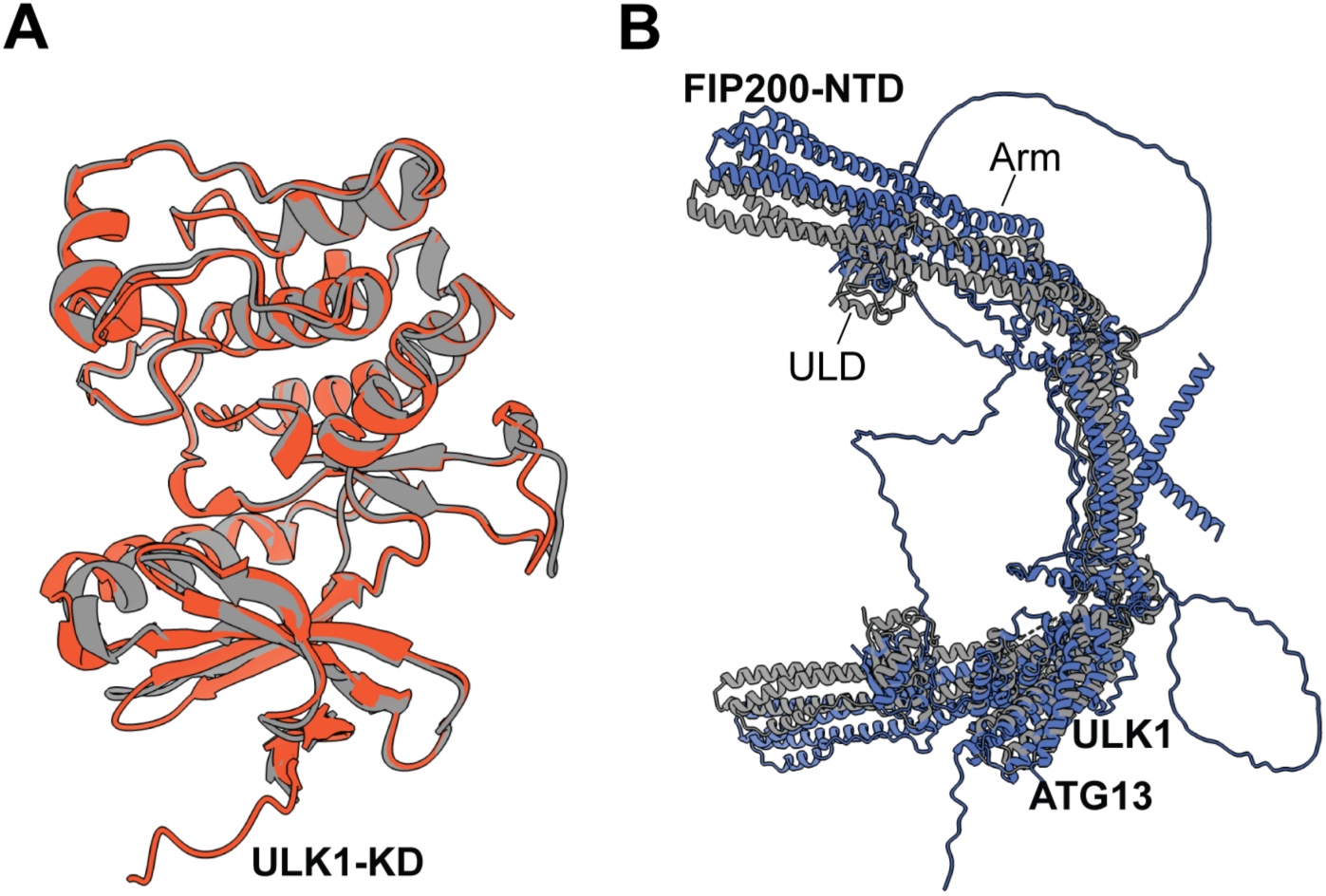
AlphaFold2 models used of ULK1-KD and ULK1C core. **(A)** Overlay of ULK1-KD AlphaFold2 prediction (orange) with PDB ID 4WNO (grey). C_a_ RMSD for experimentally resolved residues is 1.3 Å. **(B)** Overlay of ULK1C core AlphaFold2 prediction (blue) with PDB ID 8SOI (grey). C_a_ RMSD for experimentally resolved residues is 9.6 Å.

**Figure S10:**
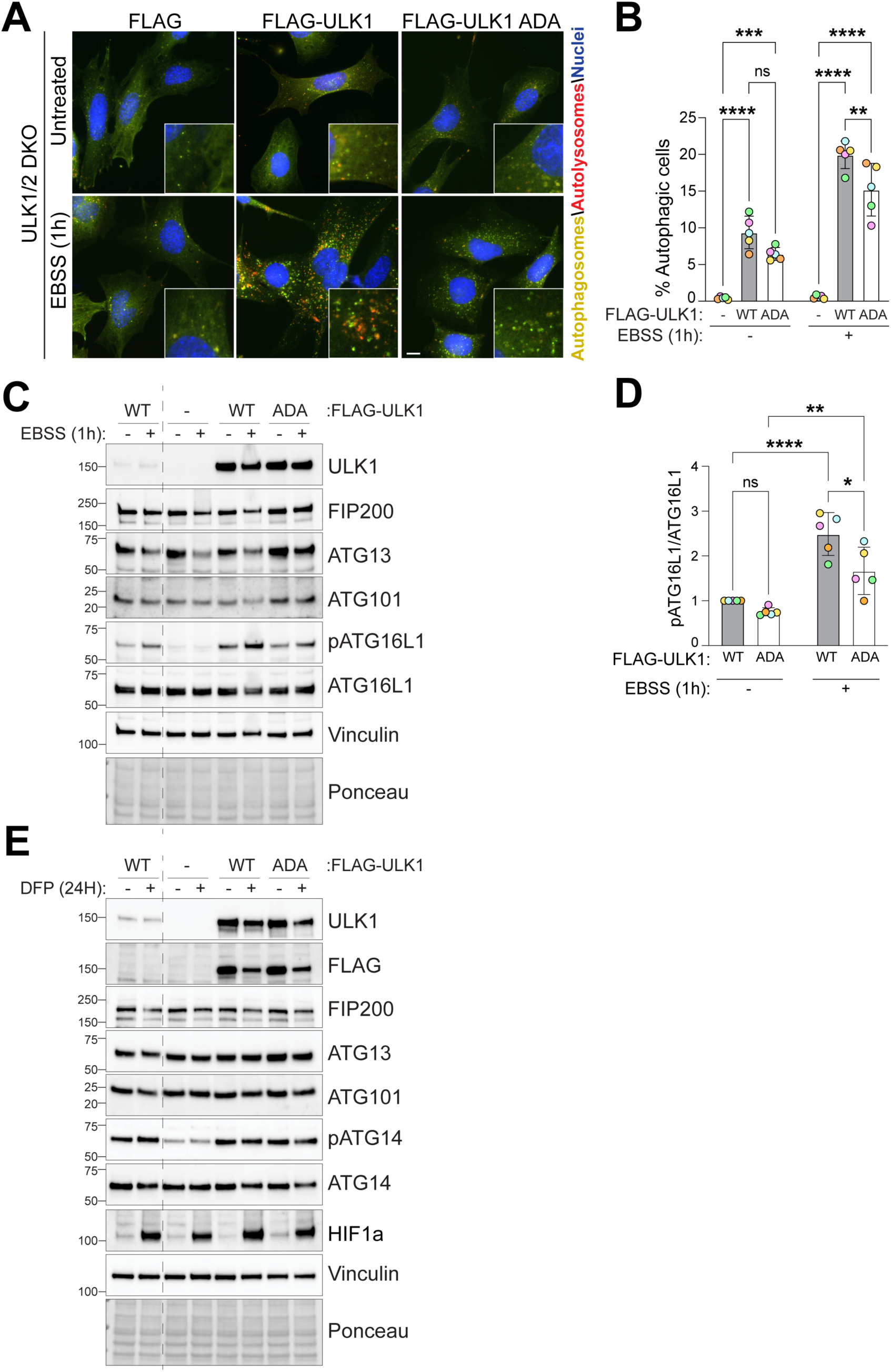
Effect on ULK1 ADA mutation on autophagy and mitophagy. (**A**) Representative wide-field images of ULK1/2 double-knockout (DKO) MEFs stably expressing an empty vector, WT, or ADA mutant FLAG-tagged ULK1 and the Auto-*QC* reporter. Cells were treated with EBSS for 1 hour prior to imaging. **(B)** Flow cytometry analysis of the mCherry/GFP ratio (n = 5 biological replicates); statistics: two-way ANOVA with Tukey’s multiple-comparisons test. **(C, D)** Representative immunoblots and quantification of the indicated proteins from cells as in (A) (n = 5 biological replicates); statistics: two-way ANOVA with Tukey’s multiple-comparisons test. (**E**) Representative immunoblots of the indicated proteins in lysates of ARPE-19 WT cells or ULK1 knock-out (KO) ARPE-19 cells stably expressing an empty vector (FLAG), WT or ADA mutant FLAG tagged ULK1. Cells were treated with 1mM DFP for 24 hours. Data information: Enlarged images are shown in the lower corners. Nuclei were stained in blue (Hoechst) and scale bars: 10 µm. All data are mean ± s.d.; Statistical significance is displayed as *P ≤ 0.05; **P ≤ 0.01; ***P ≤ 0.001; ****P ≤ 0.0001; ns, not significant.

**Table S1.**
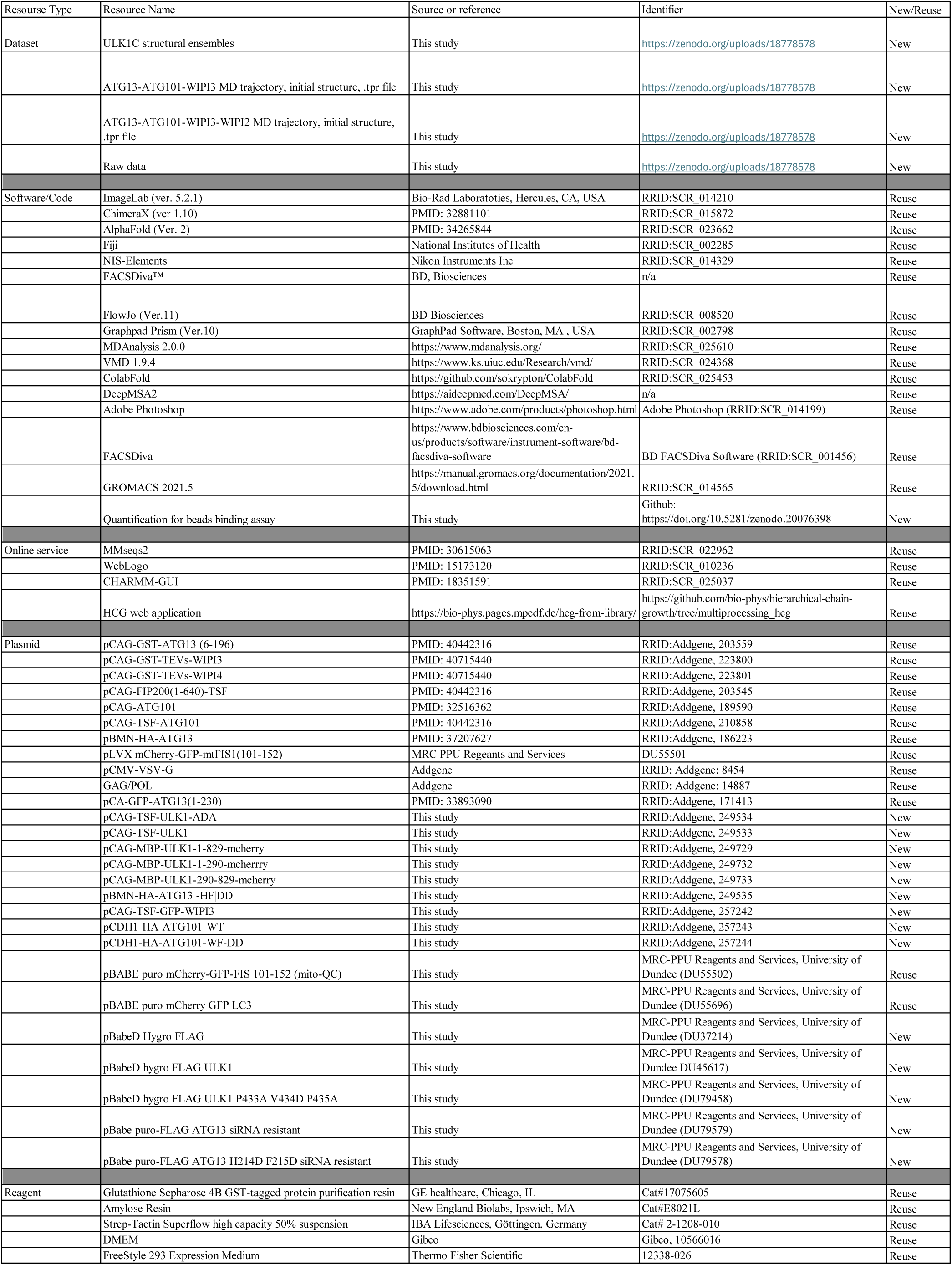
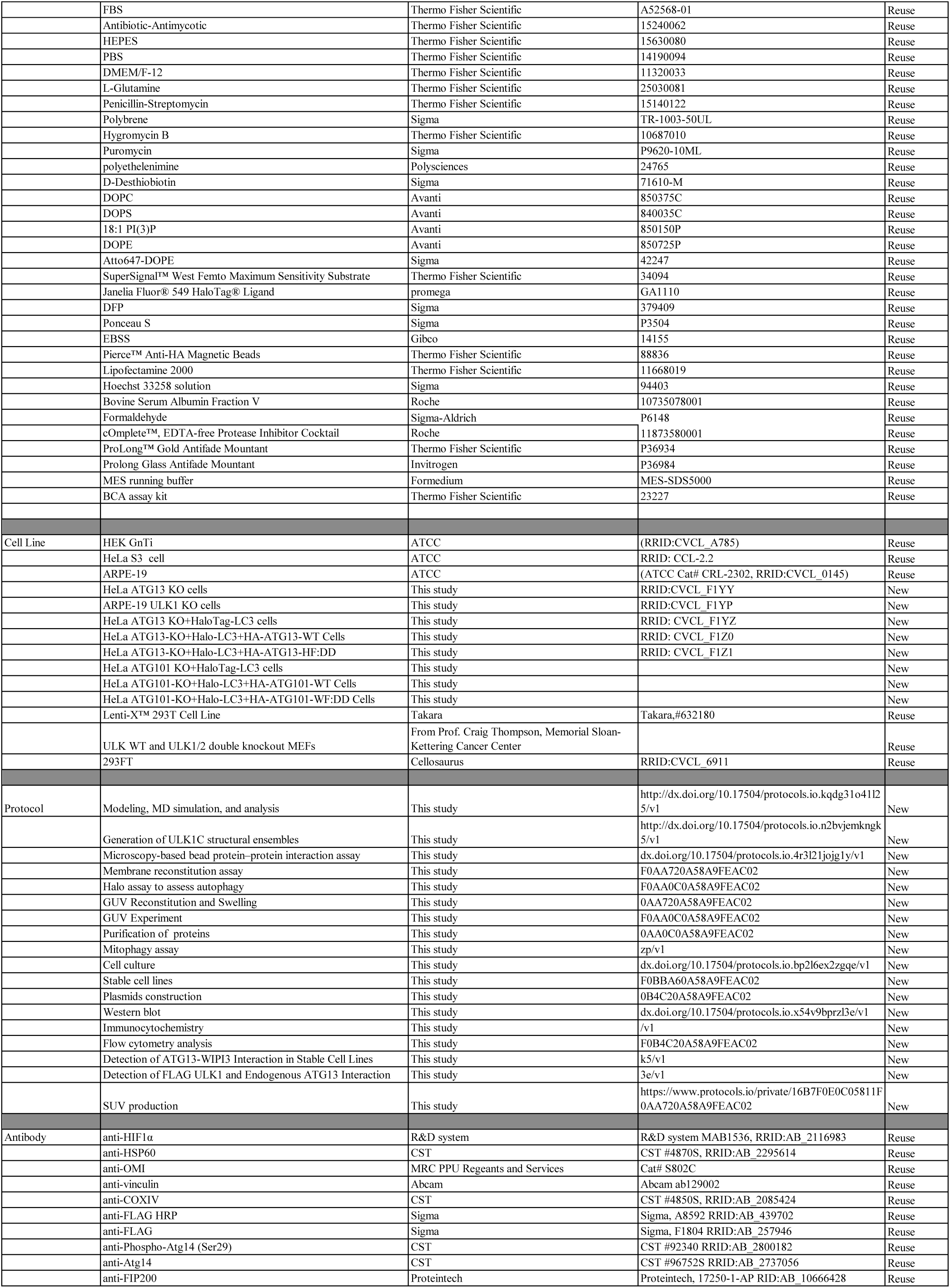
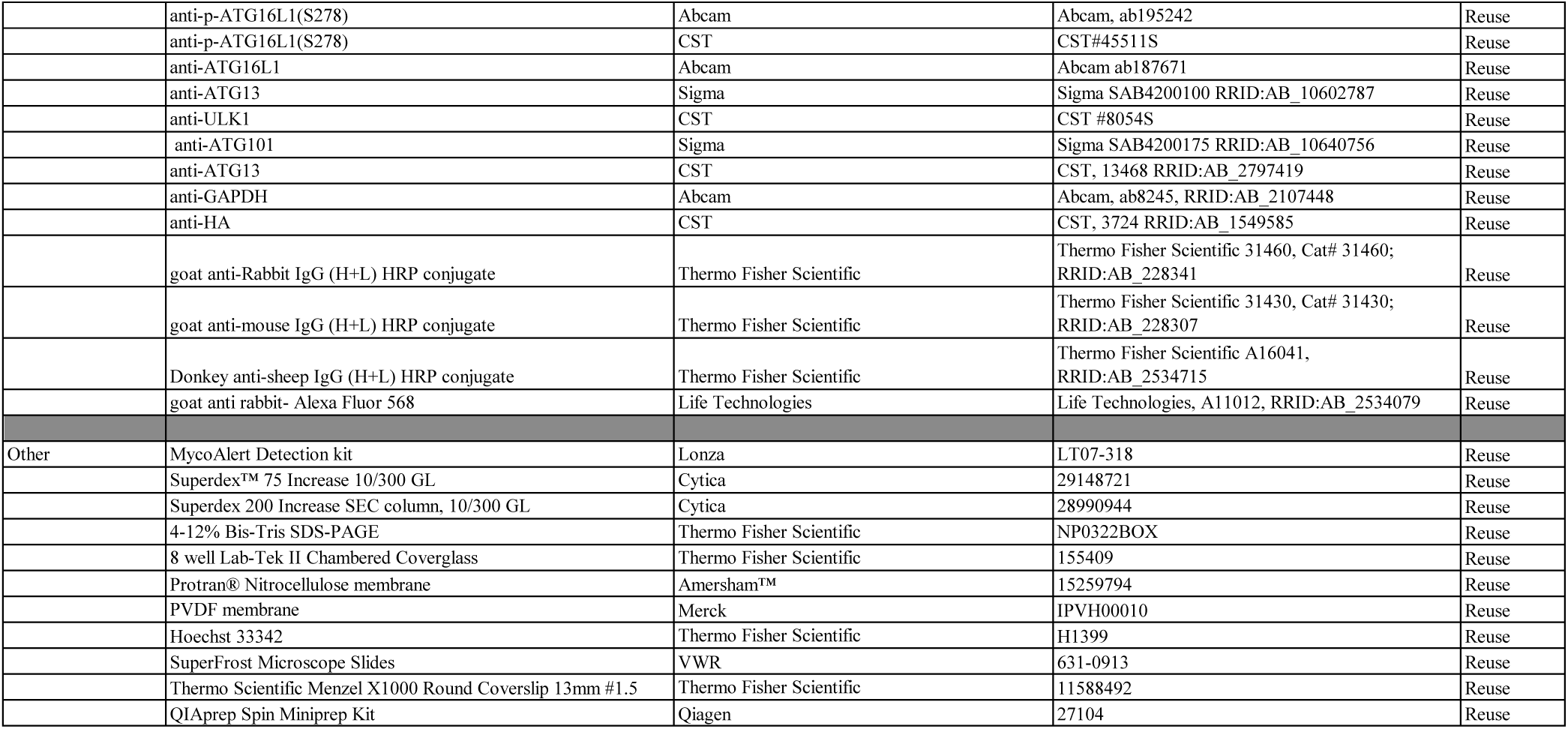

